# RNA adapts its flexibility to efficiently fold and resist unfolding

**DOI:** 10.1101/2024.05.27.595525

**Authors:** Sukjin S. Jang, Korak Kumar Ray, David G. Lynall, Kenneth L. Shepard, Colin Nuckolls, Ruben L. Gonzalez

## Abstract

Recent studies have demonstrated that the mechanisms through which biopolymers like RNA interconvert between multiple folded structures are critical for their cellular functions. A major obstacle to elucidating these mechanisms is the lack of experimental approaches that can resolve these interconversions between functionally relevant biomolecular structures. Here, we dissect the complete set of structural rearrangements executed by an ultra-stable RNA, the UUCG stem-loop, at the single-molecule level using a nano-electronic device with microsecond time resolution. We show that the stem-loop samples at least four conformations along two folding pathways leading to two distinct folded structures, only one of which has been previously observed. By modulating its flexibility, the stem-loop can adaptively select between these pathways, enabling it to both fold rapidly and resist unfolding. This paradigm of stabilization through compensatory changes in flexibility broadens our understanding of stable RNA structures and is expected to serve as a general strategy employed by all biopolymers.

## Introduction

The unique ability of biopolymers like nucleic acids and proteins to fold into and stably maintain complex three-dimensional structures is essential for their biological functions [1-5]. However, biopolymers rarely exist as a single static structure over the course of their functional lifetimes [6-8]. Instead, they form an ensemble of interconverting conformations with varying stabilities [3, 4, 9]. To understand the functions of biomolecules, it is essential to elucidate how these interconverting conformations contribute to the stability and dynamics of the ensemble as a whole.

In particular, the functional dynamics of nucleic acids, most prominently RNA, are governed by chemical properties distinct from proteins and other biomolecules. The four ribonucleotide building blocks can form stable short-range interactions in a variety of combinations that collectively yield a wide range of possible RNA secondary structures [10]. Formation of a specific secondary structure further sets the stage for establishing the longer-range tertiary interactions typically responsible for the three-dimensional organization of the structured RNA [1, 3, 11]. Thus, elucidating the conformational dynamics of these foundational RNA secondary structural elements is essential for understanding the functional dynamics of larger-scale RNA structures.

One such foundational RNA secondary structural element is a stem-loop, consisting of a double-stranded stem capped by a single-stranded loop. At least two families of stem-loops, in which the loop of each family is composed of a specific subset of four-nucleotide ‘tetraloop’ sequences (Figure 1), have been shown to be exceptionally stable and widely occurring [12-14]. Due to their prevalence and unanticipated stabilities, such ultra-stable stem-loops are thought to serve as nucleation sites for RNA folding, as elementary building blocks that anchor larger and more complex RNA structures, and as binding sites for RNA-binding proteins [15-18]. Despite decades of work investigating these ultra-stable RNA tetraloops [12, 13, 19-28], however, the unique properties of these specific tetraloop sequences that allow them to confer exceptional stability to stem-loops over other loop sequences remain unknown.

**Figure 1.**
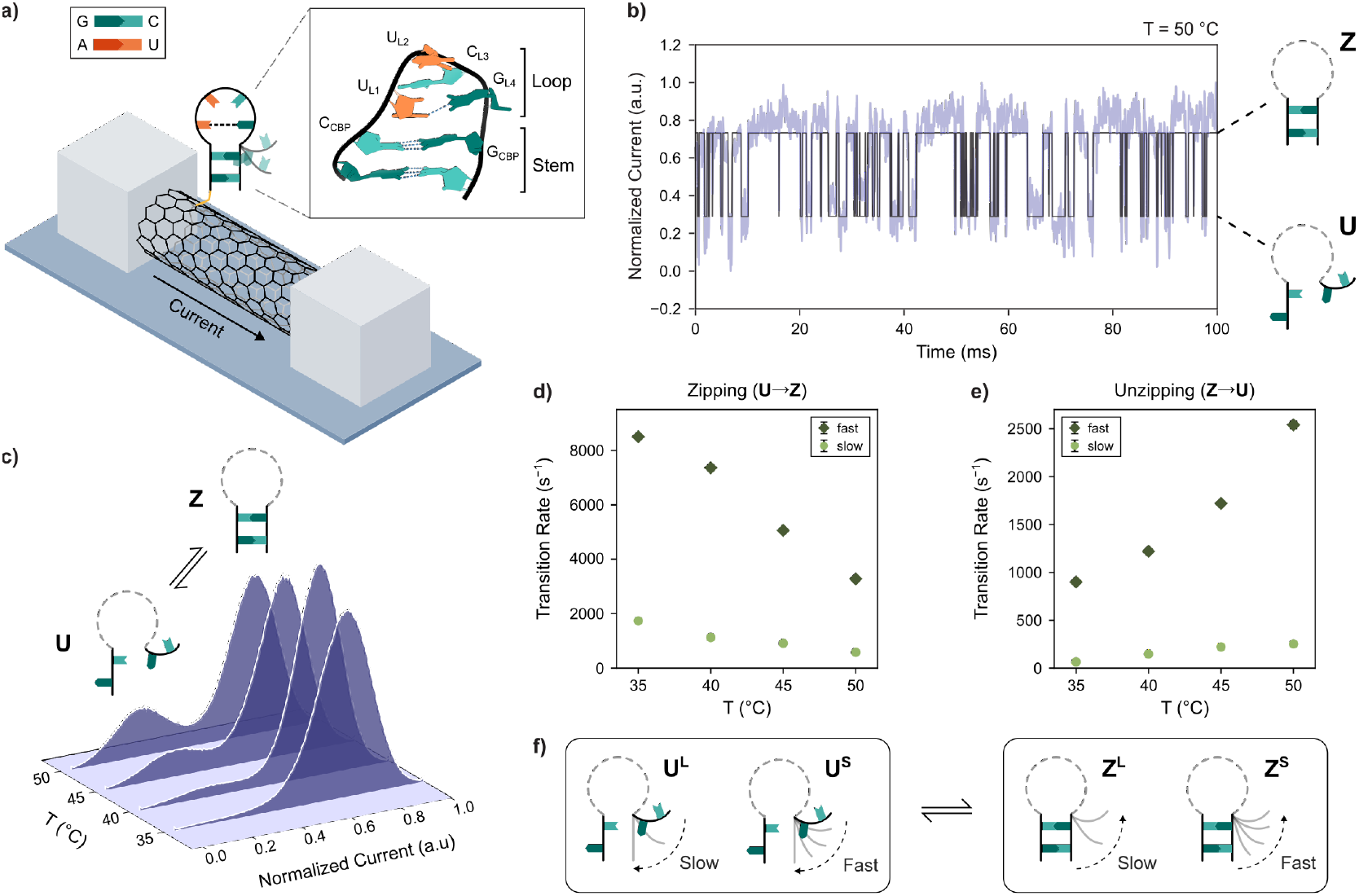
Following the zipping and unzipping kinetics of a single RNA stem-loop using smFET. **(a)** A schematic of the smFET experimental platform consisting of a single-walled carbon nanotube connected between two electrodes, with the UUCG stem-loop tethered to the carbon nanotube surface. The inset details the three-dimensional structure of the stem-loop construct (PDB ID: 1F7Y) highlighting the stem, the loop, and the nucleotides forming the loop and the closing base-pair. **(b)** A 100 ms representative observation window of a normalized current vs. time trajectory showing the real-time zipping dynamics of a single UUCG stem-loop as it transitions between **U** and **Z** conformations at 50°C. The raw signal is shown in purple and the idealized trajectory in black. **(c)** Histograms of normalized current across an entire 60 s trajectory collected at 35 °C, 40 °C, 45 °C, and 50 °C. **(d, e)** The transition rates for **(d)** zipping and **(e)** unzipping of the stem-loop at varying temperatures. **(f)** A schematic of the observed slow and fast kinetic phases, and the corresponding conformations that they transition from.

Structural studies suggest this stability arises from the ability of these tetraloops to form unique loop structures that establish a network of non-canonical interactions. For example, the ultra-stable UUCG stem-loop (containing the UUCG tetraloop sequence) has been shown to adopt a loop structure that forms a trans-G-U wobble pair between G_L4_ and U_L1_ and a stacking interaction between U_L1_ and C_L3_ (Figure 1a) [13, 28-30]. Ensemble fast kinetics studies, however, suggest that at least some of this stability arises from relatively long-lived intermediates that facilitate folding [19-21, 31]. Unfortunately, it is challenging to determine the structural identities of these intermediates from the ensemble-averaged data obtained in these studies, making it difficult to elucidate the mechanism through which these intermediates contribute to the stability. Moreover, computational studies predict that stem-loops do not exclusively fold into the single static structures captured in structural studies, but instead exist as a dynamic ensemble of closely related interconverting folded conformations [24, 25, 27, 32]. The net stability of such an ensemble would then arise collectively, from the number of populated conformations, their individual thermodynamic stabilities, and their rates of interconversion. Given these various perspectives, a single unifying model defining the exceptional stability of stem-loops has proven challenging to elucidate.

In this work, we have characterized the complete conformational ensemble of the UUCG stem-loop. Using a carbon nanotube-based nano-electronic device combined with a precise temperature-control setup, we have recorded the dynamics of a single UUCG stem-loop at microsecond resolution over a range of experimental temperatures (Figure 1). We have identified the two distinct conformations the stem-loop folds into, and resolved the differences in the loop structures adopted by these conformations, that result in varying flexibilities of the folded stem-loop. Only one of these folded conformations, the more rigid one, has been previously identified as the ‘native’ folded conformation for this motif [13, 28, 30]. We demonstrate that the non-canonical interactions in the native loop structure stabilize this native folded conformation by increasing the rate of structure formation. On the other hand, we show that when faced with destabilizing perturbations, it is the ability of the stem-loop to sample the more flexible ‘alternative’ loop structure discovered here while maintaining a zipped stem that allows it to retain its overall folded structure and resist unfolding. Beyond underpinning the physical basis for the exceptional stability of the UUCG stem-loop, this paradigm of stabilization through compensatory changes in flexibility represents a fundamental shift in our understanding of RNA folding, structure, dynamics, and function. We expect this paradigm to be a general strategy employed in the folding of other structured biopolymers. Thus, this conceptual breakthrough will be crucial in the design of mutations and small-molecule ligands to both disrupt [33] and programmatically engineer [16] specific biomolecular functions.

## Results

### The UUCG stem-loop exists as an ensemble of two zipped and two unzipped conformations

We utilized a single-molecule field effect transistor (smFET) device to investigate the dynamics of a model stem-loop construct comprising a stem of two GC base pairs and a UUCG tetraloop [20, 23, 25, 31] (Figure 1a and Table S1; see Material and Methods) over a range of temperatures. Upon tethering the stem-loop to the smFET device [34, 35] (Figures 1a and S1; see Materials and Methods), we observed a current signal that transitioned between a high-current state (normalized current of 0.7 arbitrary units (a.u.)) and a low-current state (normalized current of 0.3 a.u.) (Figure S2). Based on previous smFET studies [36-39], we could assign the 0.7 and 0.3 a.u. states to zipped and unzipped stem-loop conformations, ***Z*** and ***U***, respectively (Figure 1b). Consistent with these assignments, the population of ***U*** increased as a function of temperature, reflecting the greater propensity of the stem-loop to be unzipped at higher temperatures (Figure 1c). To confirm these assignments, we analyzed the equilibrium between these conformations using a two-state assumption (*i*.*e*., making the simplifying assumption that each state corresponded to a single conformation). This analysis accurately recapitulated the thermodynamic stability of this stem-loop as obtained from the standard two-state analysis of a conventional ensemble thermal melting experiment of the same construct (Figures S2-4 and Tables S2 and S3; see Supporting Information for a detailed description of this analysis). In addition, the populations of the two current states at each temperature were in striking agreement with the expected populations for a single stem-loop, demonstrating that the smFET device was indeed reporting on the behavior of a single, tethered stem-loop construct (Figure S4 and Table S3; see Supporting Information for a detailed discussion of this comparison). Collectively, these results validated our assignment of the two current states and demonstrated that the observed zipping dynamics originated from a single stem-loop construct that had been tethered to the smFET device.

Although the two-state thermodynamic analysis described above confirmed that our smFET experiments elicited the same stem-loop stability that is obtained from ensemble experiments, the kinetics of both the zipping and unzipping transitions in the smFET experiments could not be explained by interconversions between single zipped and unzipped conformations. Specifically, the kinetics of both the zipping and unzipping transitions consisted of at least two distinct kinetic phases at all temperatures tested (Figures 1d-e, S5 and S6, and Tables S4 and S5). The slow and fast phases of zipping transitions from ***U*** to ***Z*** arise from long- and short-lived unzipped conformations, ***U***^***L***^ and ***U***^***S***^, respectively (Figures 1f and S7a). Similarly, the slow and fast unzipping transitions from ***Z*** to ***U*** arise from long- and short-lived zipped conformations, ***Z***^***L***^ and ***Z***^***S***^, respectively (Figures 1f and S7b). Whereas the zipping transition rates decreased with increasing temperature, the unzipping transition rates increased (Figure 1d-e). This is a common characteristic of the folding kinetics of structured biomolecules [40-44].

The fact that these phases could be detected for both ***U*** and ***Z*** suggests that the rates of interconversion between ***U***^***L***^ and ***U***^***S***^ and between ***Z***^***L***^ and ***Z***^***S***^ must be slow compared to our 40 µs experimental acquisition time. The UUCG stem-loop thus exists as an ensemble of at least four conformations, two unzipped, ***U***^***L***^ and ***U***^***S***^, and two zipped, ***Z***^***L***^ and ***Z***^***S***^ (Figure 1f). However, because the normalized current values of ***U***^***L***^ and ***U***^***S***^, as well as ***Z***^***L***^ and ***Z***^***S***^, could not be distinguished, we were unable to identify transitions between the unzipped conformations or between the zipped conformations using our current analysis tools. This means we could not directly determine how these states were connected. For example, we could not ascertain whether ***U***^***L***^ transitions specifically to one of the ***Z*** conformations or to both, and the same applies for ***U***^***S***^ transitions.

### The UUCG stem-loop folds through two distinct zipping pathways

To determine the connectivity between the four observed conformations, we considered two competing kinetic schemes, one where ***U***^***L***^ predominately exchanges with ***Z***^***L***^, and ***U***^***S***^ with ***Z***^***S***^ (Figures 2a and S7c), and one where ***U***^***S***^ mainly exchanges with ***Z***^***L***^, and ***U***^***L***^ with ***Z***^***S***^ (Figures 2b and S7d). These schemes together encompass all of the possible zipping and unzipping pathways the stem-loop may take (hereafter referred to simply as zipping pathways). We then took advantage of the fact that in each kinetic scheme, the net equilibrium change in thermodynamic parameters between two arbitrary states should be the same regardless of the path used to calculate them. For instance, the net equilibrium change in enthalpy (Δ*H*_*eq*_) between two conformations (*e*.*g*., ***U***^***S***^ and ***U***^***L***^), which represents changes in stabilizing interactions, should be equal when calculated along a direct path (Path A, solid arrow in Figure 2) or an indirect path (Path B, dashed arrows in Figure 2). Similarly the corresponding equilibrium change in entropy (Δ*S*_*eq*_), which represents changes in the flexibility of the biomolecule and/or the ordering of the solvent, should be equal when calculated along Paths A and B. We calculated the individual Δ*H*_*eq*_ and Δ*S*_*eq*_ parameters between all possible pairs of conformations using our temperature-dependent measurements of zipping and unzipping kinetics (Figure S8, and Tables S6 and S7; see Materials and Methods for detailed descriptions of these calculations). We then used these parameters to calculate the net equilibrium change in thermodynamic parameters along the direct and indirect paths for the two kinetic schemes. We saw that the net changes for Paths A and B were equal only for the first scheme and not the second (Figures 2 and S7c-d, a detailed description of this comparison is provided in the Supporting Information). The zipping dynamics observed in our experiments thus overwhelmingly follow the first kinetic scheme.

**Figure 2.**
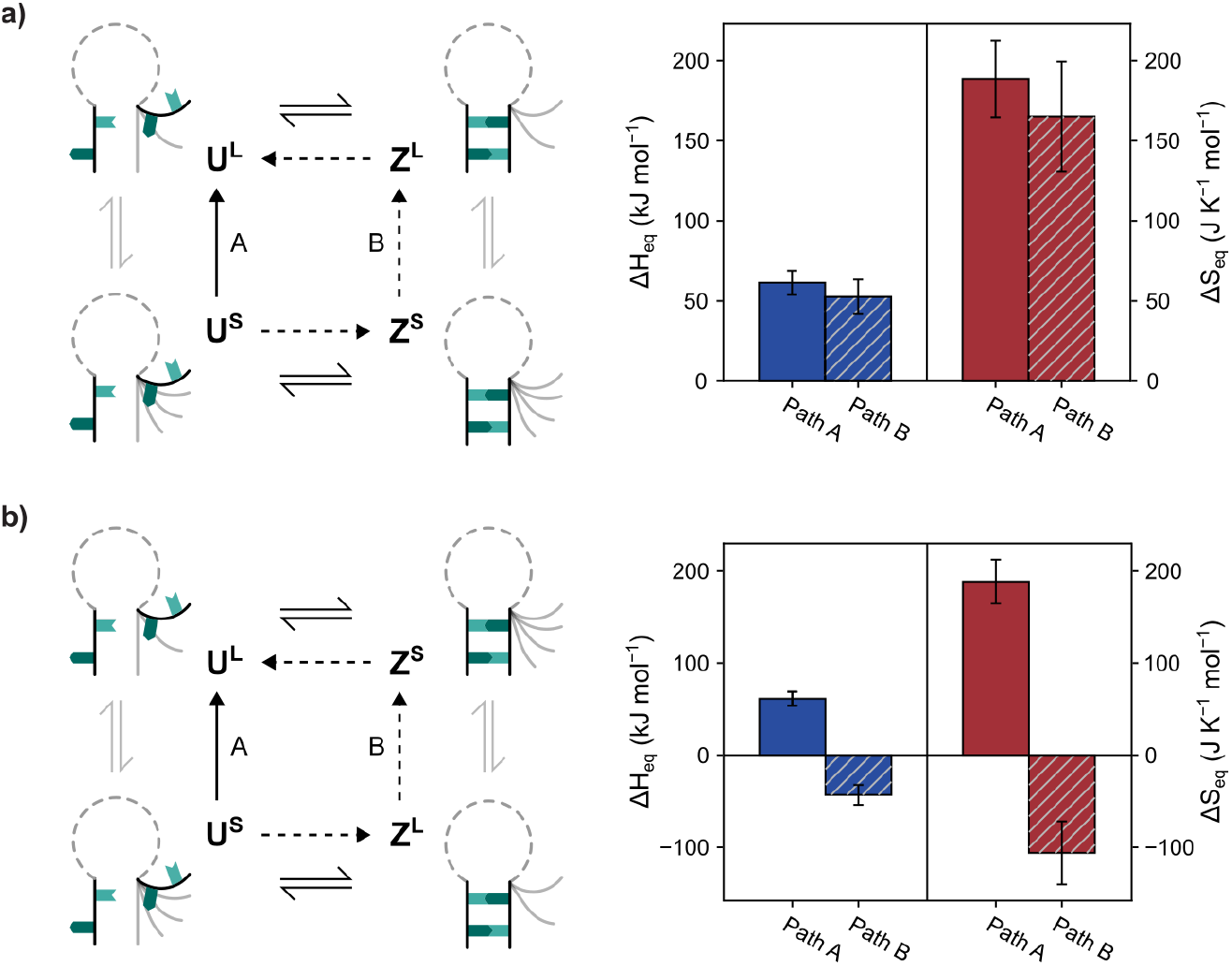
Two possible kinetic schemes for zipping and unzipping of the RNA stem-loop. The hypothetical kinetic schemes (left) showing predominant transitions **(a)** between **U**^**L**^ and **Z**^**L**^, and **U**^**S**^ and **Z**^**S**^, and **(b)** between **U**^**L**^ and **Z**^**S**^, and **U**^**S**^ and **Z**^**L**^, and the corresponding thermodynamic cycle depicting the direct Path A (solid arrow) and the indirect Path B (dashed arrows) between **U**^**S**^ and **U**^**L**^. The undetected transitions between the **U** conformations and **Z** conformations are shown with grey equilibrium arrows. The bar graphs (right) depict the calculated **ΔH**_**eq**_s (blue) and **ΔS**_**eq**_s (red) for these paths for both **(a)** and **(b)**. The error bars represent the propagated errors of fit for the linear fits of free energies as a function of temperature (Figure S8).

UUCG stem-loop zipping is therefore characterized by two distinct kinetic pathways that exist in parallel to one another, a slow pathway consisting of transitions between ***U***^***L***^ and ***Z***^***L***^, and a fast pathway consisting of transitions between ***U***^***S***^ and ***Z***^***S***^ (Figure 2a). In both pathways, zipping transitions occur at a faster rate than unzipping transitions. Thus, within the temperature range of our experiments, which is below the stem-loop melting temperature, *T*_*m*_, of 57.0 °C, the equilibria for both pathways favor the zipped conformations over the corresponding unzipped conformations. Notably, upon an increase in temperature, the slow pathway becomes progressively more populated over the fast pathway (Tables S4 and S5), revealing that this zipping pathway of transitions between ***U***^***L***^ and ***Z***^***L***^ is more dominant at higher temperatures.

### The two zipping pathways arise from differences in the structure of the loop

We next sought to identify what RNA structures correspond to ***U***^***L***^, ***U***^***S***^, ***Z***^***L***^, and ***Z***^***S***^. We expected our experimentally determined *ΔH*_*eq*_ s (Figure S8) to predominantly report on the relative stabilizing interactions present in these conformations, and the *ΔS*_*eq*_ s (Figure S8) to predominantly report on their conformational flexibilities. ***U***^***L***^, in which the stem-loop possesses the least stabilizing interactions and is the most flexible, thus corresponds to the fully unfolded structure (Figure 3a). Analogously, ***Z***^***S***^, in which the stem-loop forms the most stabilizing interactions while also being the most rigid, corresponds to the previously characterized fully folded, native stem-loop structure (Figure 3a).

**Figure 3.**
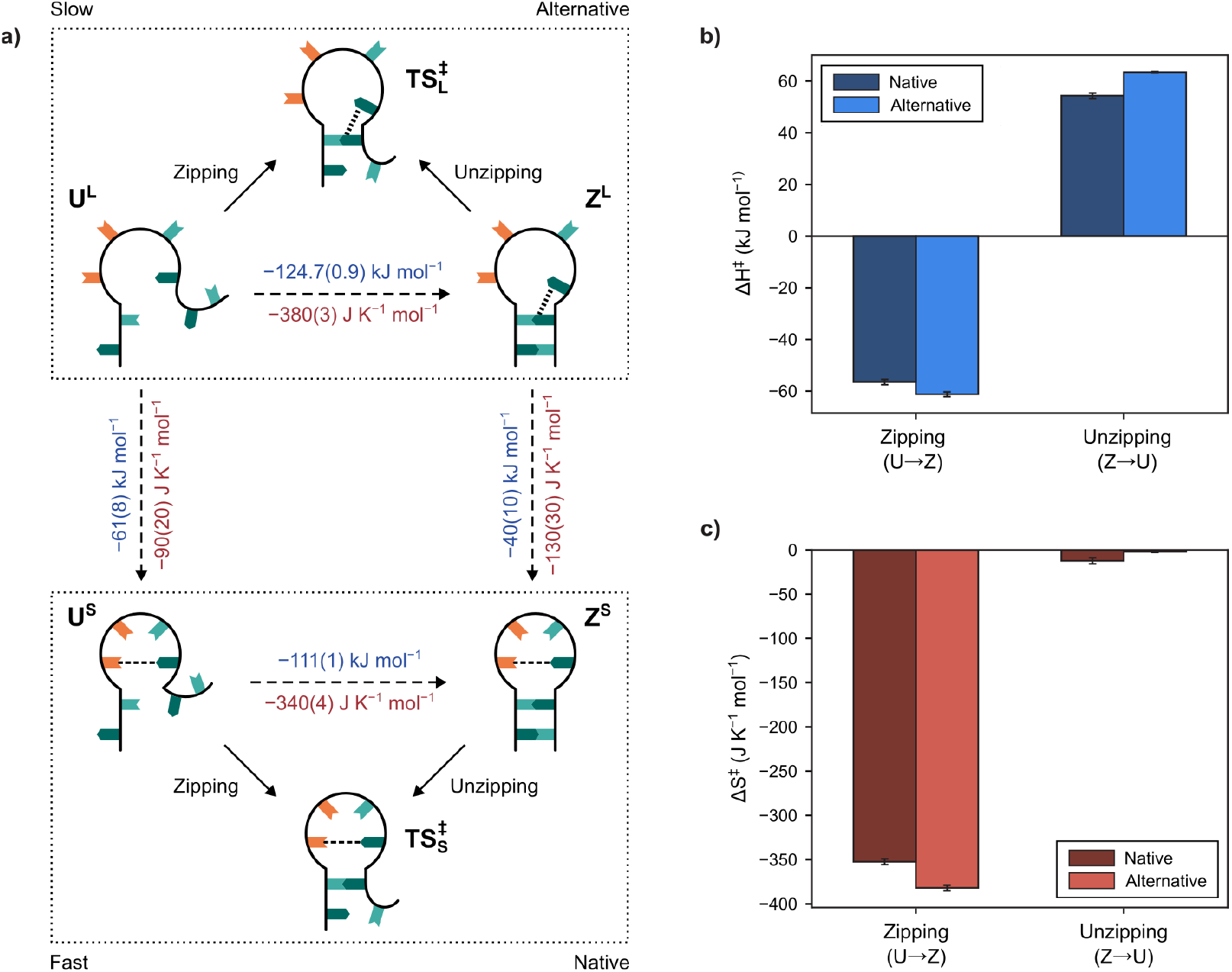
The structural identities of the U and Z conformations. **(a)** Schematic represenations of the unzipped conformations (**U**), transition states (**TS**^‡^), and zipped conformations (**Z**) along the alternative (above) and native (below) zipping pathways. The calculated **ΔH**_**eq**_s (blue) and **ΔS**_**eq**_s (red) between each of the observed **U** and **Z** conformations are shown. **(B)** Activation enthalpies (**ΔH**^‡^s) for the native (dark blue) and alternative (light blue) pathways. **(C)** Activation entropies (**ΔS**^‡^s) for the native (dark red) and alternative (light red) pathways. The errors (in parentheses) and error bars represent the propagated errors of fit for the linear fits of free energies as a function of temperature (Figure S8).

To determine the identity of ***Z***^***L***^, we compared the two zipped conformations. The stem-loop has less stabilizing interactions and is more flexible in ***Z***^***L***^ than in ***Z***^***S***^ (Figure 3a). Since ***Z***^***L***^ and ***Z***^***S***^ both correspond to zipped conformations, the RNA must have similar stem structures in both; it is therefore the structure of the UUCG loop that differs between the two conformations. Having established that this loop adopts the previously characterized native structure in ***Z***^***S***^ [28, 30], the loop in ***Z***^***L***^ must adopt an alternatively folded structure that lacks some of the native ***Z***^***S***^ stabilizing interactions (*e*.*g*., the *trans*-G_L4_-U_L1_ wobble pair and/or the stacking interaction between U_L1_ and C_L3_) and is therefore more flexible. The absence of both of these interactions easily accounts for the ∼40 kJ mol^−1^ difference in stabilizing interactions that we observe between the two ***Z*** conformations. While the UUCG loop has previously been predicted to be capable of adopting multiple non-native structures [25, 45, 46], this alternatively folded loop structure has not previously been experimentally observed.

To determine the identity of ***U***^***S***^, we similarly compared the two unzipped conformations. ***U***^***S***^ possesses more stabilizing interactions and is significantly more constrained than ***U***^***L***^ (Figure 3a). Since ***U***^***S***^and ***U***^***L***^ both represent unzipped conformations, these differences must also arise from differences in the structures of the loop in ***U***^***S***^ and ***U***^***L***^. Since ***U***^***S***^ directly transitions to ***Z***^***S***^, we hypothesize that ***U***^***S***^ corresponds to a partially unfolded, frayed structure in which the stem-loop is unzipped, but the loop retains its natively folded structure (Figure 3a). Persistence of the native loop structure in ***U***^***S***^ is in good agreement with the ∼61 kJ mol^−1^ difference in stabilizing interactions we observe between ***U***^***S***^ and ***U***^***L***^. Our results therefore reveal a hierarchical mechanism of UUCG stem-loop folding in which the loop structure dictates whether the stem-loop folds along a ‘native’ zipping pathway from ***U***^***S***^ to the natively folded ***Z***^***S***^ or an ‘alternative’ zipping pathway from ***U***^***L***^ to the alternatively folded ***Z***^***L***^.

### The alternative UUCG loop structure forms base-stacking interactions with the zipped stem

We next sought to further characterize the alternative stem-loop structure we discovered by identifying stabilizing interactions in ***Z***^***L***^ that are not present in ***Z***^***S***^. Unzipping along the alternative pathway resulted in a greater disruption of interactions than unzipping along the native pathway (Figure 3a). This greater disruption in stabilizing interactions suggested that the stem itself is more stable and resistant to unzipping in the alternative pathway than in the native pathway. We stress that this increased stability refers to the stem alone; when considering the stabilizing interactions contributed by both the stem and the loop, the natively folded ***Z***^***S***^ is more stable overall than the alternatively folded ***Z***^***L***^. Nonetheless, this greater stability of the stem in ***Z***^***L***^ relative to ***Z***^***S***^ is counterintuitive based on the widely described role of the native UUCG loop structure in preferentially stabilizing the zipped stem-loop [13, 26, 28, 29].

Since stem unzipping is not expected to lead to any changes in the loop structure itself, the increased stability of the zipped stem in ***Z***^***L***^ must arise from the alternative loop structure forming more interactions with the stem than the native loop structure. Previous computational studies guided our identification of these interactions [25]. In the native loop structure, G_L4_ in the UUCG loop (Figure 1a) adopts the *syn*-configuration to form the *trans*-G-U wobble base pair. An *anti*-configuration of G_L4_ would inhibit the formation of this *trans*-G-U wobble interaction, but instead allow G_L4_ to stack against the adjacent G_CBP_ (G in the closing base pair; Figure 1a) [47]. We hypothesize that it is this stacking interaction in ***Z***^***L***^ that stabilizes the zipped stem and needs to be overcome during unzipping in the alternative pathway (Figure 3a). Intriguingly, the stronger stabilizing interactions that the stem makes with the alternative loop structure *versus* the native loop structure, suggests that the role of the native loop structure in stem-loop folding goes beyond the formation of these interactions.

### Formation of the closing base-pair determines the rate of stem zipping

To explore the role of the native loop structure in stem-loop folding, we sought to identify the transition state (***TS***^‡^) RNA conformations that comprise the barriers to the zipping and unzipping transitions. We used our temperature-dependent measurements of zipping to describe these barriers in terms of changes in stabilizing interactions (*i*.*e*., activation enthalpy, Δ*H*^‡^s) and in conformational flexibility (*i*.*e*., activation entropy, Δ*S*^‡^s) during the rate-determining steps of these transitions [48]. For zipping transitions in both pathways, the negative Δ*S*^‡^s indicate that the ***TS***^‡^s are significantly more constrained than the corresponding unzipped conformations (Figures 3b and 3c). Thus, these barriers are dominated by the challenge of constraining the stem in order to zip. In contrast, the ***TS***^‡^ s are almost equally as constrained as the corresponding zipped conformations, but contain less stabilizing interactions (Figures 3b and 3c). Specifically, these ***TS***^‡^s both possess half of the stabilizing interactions that are required for the stem to fully zip (Figures 3a and 3b, and Table S6). Given that zipping of the stem in our construct involves formation of only two base-pairs, this result suggests that one, and only one, of these base-pairs is already formed in the ***TS***^‡^s, and it is the formation of this base-pair, associated with a significant loss in flexibility of the RNA, that determines the rate of stem zipping in both pathways.

Comparison of the two pathways revealed that formation of the first base-pairing interaction leads to greater stabilization (*i*.*e*., a larger Δ*H*^‡^) in 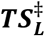 than in 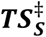 (Figure 3b). We attribute this increased stabilization to the additional base-stacking interaction formed between G_L4_ and G_CBP_ in 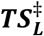 Since this stacking interaction is likely to be formed only when G_CBP_ is base-paired, it is specifically the closing base pair that is formed in 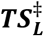. Given the overall similarity between zipping in the alternative and native pathways, we expect 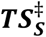 to also contain just the closing base pair. The formation of the second base pair also leads to greater stabilization between 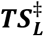 and ***Z***^***L***^, than between 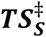 and ***Z***^***S***^ (Figure 3b), as would be expected when the stacking network in 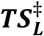 subsequently expands to the outer base pairs in ***Z***^***L***^.

Our results thus reveal that it is the loss of flexibility associated with formation of the closing base-pair of the stem that serves as the barrier for zipping. On the other hand, the barrier for unzipping is almost exclusively due to the disruption of the outer base-pair (Figure 3a), and therefore, is not associated with any significant increase in flexibility (Figure 3c). The small differences in entropy that we observe between the ***Z*** conformations and the ***TS***^‡^s (*i*.*e*. the Δ*S*^‡^s for unzipping) likely arise from changes in the ordering of the solvent when the outer base pair forms [49] rather than from changes in the flexibility of stem-loop itself.

### The native loop structure facilitates zipping by increasing the proximity between the closing bases

The results described above suggest that any RNA structure that increases the proximity between the two closing bases would facilitate zipping by reducing the change in flexibility that is needed to form the stem. This is the case for ***U***^***S***^, where formation of the first base-pair is associated with a smaller loss of flexibility than for the fully unfolded ***U***^***L***^ (Figure 3c). The pre-organization of the native loop structure in ***U***^***S***^ thus leads to a faster rate of zipping from ***U***^***S***^ (Figure 1d). Counterintuitively, in the alternative pathway, the stacking interaction between G_L4_ and G_CBP_ leads to a further loss of flexibility that needs to be overcome to form the stem, resulting in a slower rate of zipping despite the greater stabilization that results from forming this interaction.

A powerful way to visualize how the native loop structure facilitates stem-loop folding is through a schematic of the potential energy landscape for the folding mechanism of the UUCG stem-loop (Figure 4a). In this hierarchical landscape, two large basins represent the ensembles of conformations in which the UUCG tetraloop adopts the native or alternative loop structure. These basins each contain one low-potential-energy well corresponding to ***Z***^***L***^ or ***Z***^***S***^, while regions of the basins outside each well correspond to ***U***^***L***^ or ***U***^***S***^. The depths of the basins and wells represent the relative stabilities due to the interactions present in the corresponding conformations, while their widths represent their relative flexibilities. The natively folded ***Z***^***S***^, which has more stabilizing interactions and is more constrained than ***Z***^***L***^, corresponds to a deeper and narrower well. Since our existing data analysis tools cannot identify the transitions between the two basins in our experimental data (*i*.*e*., the transitions between the slow and fast phases of zipping and unzipping in our current trajectories), we represent the barrier between the two basins as a discontinuity.

**Figure 4.**
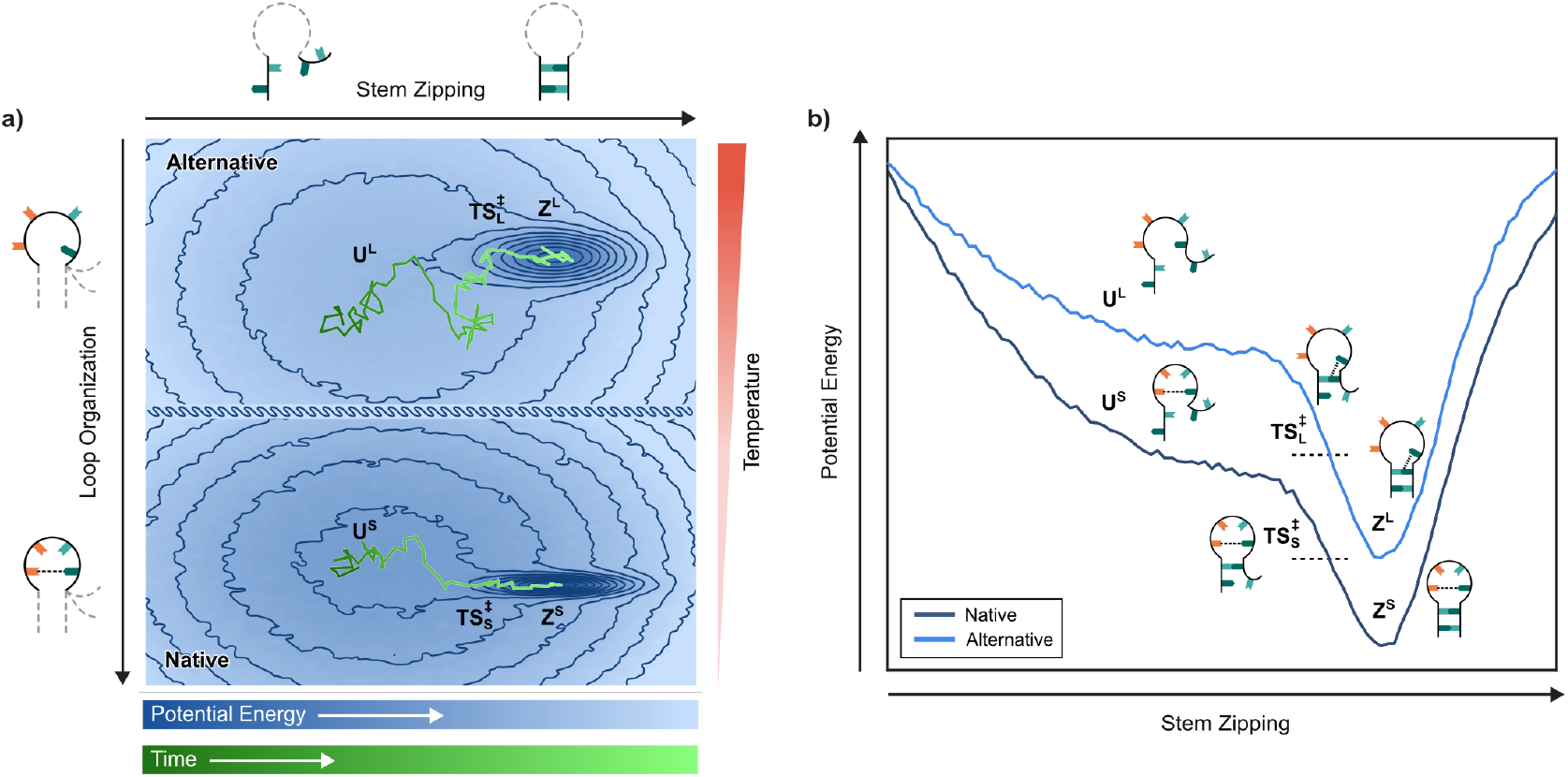
The physical basis of UUCG stem-loop stability. **(a)** The hypothetical hierarchical potential energy landscape for the folding of the stem-loop along the two reaction coordinates of loop organization and stem zipping. Each point in the landscape represents the potential energy of a specific stem-loop conformation projected onto these coordinates. The border between the native and alternative loop structure basin is shown as discontinuous to represent the lack of information about the barriers between the two in our analysis of the data. Two hypothetical tracks representing stem-loop folding in both basins are shown, a short track capturing the fast guided search for **Z**^**S**^ in the narrow native loop structure basin, and a more difussive track representing the slower search for **Z**^**L**^ in the wider alternative loop structure basin. As temperature is increased, the alternative loop structure basin is preferred. **(b)** A hypothetical representation of the one-dimensional potential energy along the stem zipping reaction coordinate for the native and alternative pathways. The positions of **U**^**L**^, **Z**^**L**^, and 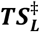 in the alternative pathway, and **U**^**S**^, **Z**^**S**^, and 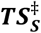 in the native pathway, are shown for both **(a)** and **(b)**.

The process of stem-loop folding from the fully unfolded ***U***^***L***^ to natively folded ***Z***^***S***^ is the conformational search on this landscape from the shallow and wide non-native loop basin into the deep and narrow ***Z***^***S***^ well. The UUCG stem-loop has evolved to pay the cost associated with the loss of flexibility for this transition in two steps, using a set of hierarchical loop and stem dynamics. In the first step, part of this cost is paid by organizing the loop into the native structure, forming a metastable intermediate, ***U***^***S***^, that nucleates the process of stem-loop folding, and shifts to the deeper and, in particular, narrower native loop basin. This constrains the conformational search, allowing transitions to the ***Z***^***S***^ well from ***U***^***S***^ to be faster (as shown by the shorter track in Figure 4a). The rate-limiting step for zipping of the stem *via* this pathway is the second step, requiring loss of the remaining flexibility in the RNA to reach 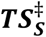, which contains only the closing base-pair and is situated at the entrance to the ***Z***^***S***^ well. The downhill pathway from ***U***^***L***^, through ***U***^***S***^, to ***Z***^***S***^ allows the loss of flexibility at each of the two steps to be coupled to a gain in stabilizing interactions (Figure 4b).

In summary, the ability of the UUCG sequence to form the native loop structure in the absence of a stem facilitates folding of the stem-loop into ***Z***^***S***^ in two ways. First, it creates a hierarchical separation between loop organization and stem zipping such that the entropic cost of stem-loop folding can be paid over two steps, resulting in a guided conformational search for the natively folded structure of the stem-loop. Second, by pre-organizing the loop structure to nucleate folding in the first step, formation of the closing base-pair in 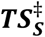 is made more probable, cooperatively facilitating the second step of stem formation. Taken together, this describes the mechanism through which pre-organization of the tetraloop significantly reduces the conformational space of the stem-loop and expedites the folding process.

### The alternative zipped conformation resists unfolding through compensatory changes in flexibility

In addition to the guided-search mechanism of folding along the native pathway described above, we see that the alternative folding pathway also contributes significantly to the observed stability of the UUCG stem-loop. This is specifically observed under conditions, such as higher temperatures, where the alternative zipping pathway becomes progressively more dominant (Tables S4 and S5). This effect can also be visualized using the potential energy landscape for UUCG stem-loop folding (Figure 4a). The basin for the alternative zipping pathway is very similar to the native one described above, with a ***Z***^***L***^ well, and a transition state, 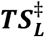, at the entrance of the well (Figure 4a). Importantly, however, the ***Z***^***L***^ well is wider than the ***Z***^***S***^ well. When faced with destabilizing perturbations like higher temperatures, the conformational search for this well from the fully unfolded ***U***^***L***^ is easier than for the narrower ***Z***^***S***^ well. This is represented in the potential energy landscape by the more diffusive track in the alternative pathway basin—representing more thermal fluctuations of the RNA—still being able to find and zip into the wider ***Z***^***L***^ well (Figure 4a). Secondly, the wider ***Z***^***L***^ well also reflects how the stem-loop can adopt a larger variation of more flexible loop structures while maintaining a zipped stem, which is crucial for remaining folded at higher temperatures. Finally, due to the additional base stacking interaction between the stem and the loop, ***Z***^***L***^ possesses a higher barrier to exit than ***Z***^***S***^, and once in the well, transitioning out is less probable (Figure 4b). Altogether, this shows how the stem-loop can conditionally favour more flexible RNA conformations by shifting its ensemble to the more flexible ***Z***^***L***^ in order to resist unfolding at higher temperatures.

***Z***^***L***^ is clearly a kinetic trap in the folding process of the native UUCG stem-loop structure. However, the crucial role of this conformation in the stability of the folded stem-loop structure as a whole leads us to suggest that ***Z***^***L***^ does not represent merely a misfolded conformation, but is instead a previously unidentified, functionally relevant alternative structure of this RNA. While beyond the scope of this work, additional functional roles for this alternative UUCG loop structure need to be further explored. We note, for instance, that the alternative loop structure in ***Z***^***L***^ (see above, Figure 3a) may position the UUCG stem-loop to make tertiary contacts with other RNAs or proteins by freeing the bases in the loop. This previously uncharacterized conformation may thus have unexplored biological roles beyond stabilizing the stem-loop by resisting unfolding.

## Discussion

We have elucidated the complete conformational ensemble of a fundamental RNA structural motif, the UUCG stem-loop, using a high-time-resolution, single-molecule experimental approach uniquely suited for studying the dynamics of such biopolymers. We characterize, in molecular detail, the competing pathways, metastable intermediates, and kinetic traps that collaborate to give rise to the exceptional thermodynamic stability of this RNA stem-loop. Our findings reveal that the UUCG stem-loop uses a hierarchical conformational search process to guide folding into the fully folded native structure. We further show how the specific ordering of this hierarchy into the two steps of loop structure organization and stem zipping facilitates the efficient collapse of the flexible RNA into the constrained native folded state (Figure 4a) in the native zipping pathway—an entirely entropic effect. We further discover how this RNA stem-loop resists unfolding when challenged with destabilizing perturbations by shifting its equilibrium to a more flexible alternatively folded conformation (Figure 4a), a conformation that may also have additional, hitherto uncharacterized biological roles. Our results elucidate how these dual mechanisms of constrained conformational search and compensatory changes to flexibility collaborate to give rise to the exceptional thermodynamic stability of the UUCG stem-loop.

The folding mechanism we have elucidated here has clear implications for our views of the biological roles of UNCG stem-loops (where N denotes any nucleotide). UNCG stem-loops, including UUCG stem-loops, are typically considered to only serve as ultra-stable structural scaffolds [15]. This perspective has predominately been informed by static structures of the fully folded, native structure of the UUCG stem-loop, which exhibits only a limited set of tertiary interactions [13, 26, 28, 29]. However, our results show that this native structure is not the sole folded structure that can be populated by this stem-loop and, indeed, is the less populated structure under certain conditions. In contrast to the native loop structure, the alternative loop structure we observe here is expected to allow for greater tertiary interactions. Our results thus suggest that the UUCG stem-loop may play a biological role in interacting with other biomolecules through this alternative structure. Our hypothesis is supported by a recent study analyzing existing structures of RNA stem-loops, which found that UNCG tetraloops can populate multiple conformations referred to as ‘non-native’ when bound to other biomolecules [50]. In particular, the ability of the UUCG stem-loop to shift between the native and alternative loop structures in response to perturbations sets up an enticing model for the regulation of stem-loop interactions with other biomolecules. In this model, perturbations to the UUCG stem-loop, whether through changing conditions or the allosteric effect of the binding of ligands or other biomolecules, can shift the equilibrium between the two loop structures, tuning the ability of the stem-loop to interact with partners. This, of course, is a hypothesis which needs to be further investigated in light of our findings.

Finally, the complete folding mechanism of the UUCG stem-loop that we reveal here (Figure 4) is in striking contrast to our current understanding, which attributes the exceptional thermodynamic stability of this RNA motif solely to the interactions within the unique native UUCG loop structure [13, 28-30]. While beyond the scope of this work, it will be worthwhile to determine how this folding mechanism maps onto other thermodynamically stable RNA motifs, including GNRA stem-loops (where N denotes any nucleotide and R any purine nucleotide), which are thought to have evolved a strategy for achieving their exceptional thermodynamic stability that is independent and distinct from that of the UNCG stem-loops [51]. It is worth noting, in this context, that the strategies for thermodynamic stability we describe here do not appear to be specific to stem-loops—or indeed, to RNA—and thus, we expect to see them in the folding mechanisms of a wide range of biomolecules. In particular, the strategy we have discovered here of resisting destabilizing perturbations through compensatory changes in the flexibility of the biomolecule represents an exciting addition to our understanding of biomolecular stability, the implications of which must be explored for broader classes of structured biopolymers.

## Materials and Methods

### Design of the smFET experiments

The single-walled carbon nanotube (SWCNT)-based smFET experimental platform, including details describing the theoretical basis for the function of an smFET device, practical aspects of how the current is measured, and how the temperature is controlled, has been previously described [34, 38, 52]. Briefly, an smFET device provides a measure of current through the SWCNT as a function of time. This current signal is extremely sensitive to the local charge environment near the surface of the SWCNT where the biomolecule is tethered. Because conformational changes, including folding and unfolding, of a biomolecule such as an RNA alters the electrostatic charge distribution near the surface of the SWCNT, the current *vs*. time trajectory (hereafter referred to as the current trajectory) recorded by an smFET device very sensitively reports on these conformational changes.

The smFET experiments reported in this work utilized a chemically synthesized RNA stem-loop construct (Integrated DNA Technologies) that we refer to here as h-SL-bio (Table S1). h-SL-bio consisted of a stem containing two G-C base-pairs and a loop composed of a four-nucleotide UUCG ‘tetraloop’ sequence. The 5’ end of h-SL-bio was modified with a hydrazide functional group that allowed us to tether the construct to the surface of the SWCNT of our smFET devices. The 3’ end of the h-SL-bio was further functionalized with a biotin for potential downstream validation of tethering with additional markers. Fortunately, tethering of h-SL-bio to the surface of SWCNT could be directly validated using the current trajectories recorded by the smFET device (see below and Figures S1 and S2) and therefore downstream validation of tethering using the biotin functional group was unnecessary. To confirm that the 3’ biotin moiety does not alter the overall thermodynamic stability of the UUCG stem-loop, we conducted additional thermal melting experiments in which we compared the h-SL-bio construct to the same chemically synthesized RNA construct (Integrated DNA Technologies) that did not contain the biotin modification (a construct that we refer to as h-SL, Table S1). The presence of the biotin did not perturb the thermodynamic properties of the stem-loop, demonstrating that it did not alter the folding dynamics of the stem-loop (Figure S3 and Table S2).

A single molecule of the h-SL-bio construct was tethered to the SWCNT surface using an electrically coupled diazonium reaction [34, 35] (Figure S1a). Briefly, a source-drain current *vs*. liquid-gate voltage (*I*_*sd*_ -*V*_*lg*_) curve was obtained from the smFET device prior to running the diazonium reaction by applying a constant source-drain voltage (*V*_*sd*_) and a sweeping *V*_*lg*_ from −0.5 V to +0.5 V through the platinum electrode of the smFET device in Recording Buffer (10 mM sodium phosphate (Na_2_HPO_4_)/sodium dihydrogen phosphate (NaH_2_PO_4_), pH = 7.0). Before the introduction of 4-formylbenzene diazonium hexafluorophosphate (FBDP) to the devices, the device was placed at a *V*_*lg*_ of −0.5 V to prevent spurious diazonium reactions. The device was subsequently incubated in 300 µM FBDP dissolved in Recording Buffer to initiate the reaction. All devices were then slowly ramped towards a positive *V*_*lg*_ until a clear downward transition in current was observed, indicating the coupling of FBDP to the SWCNT surface (Figure S1b). Following the FBDP reaction, all devices were rinsed with 1 mL of Recording Buffer at 50 °C to remove any residual FBDP salts, and a new *I*_*sd*_-*V*_*lg*_ curve was obtained with the same conditions as before the diazonium reaction. The comparison between the *I*_*sd*_-*V*_*lg*_ curves shows a clear reduction in conductance before and after FBDP exposure (Figure S1c). To tether each stem-loop construct, the hydrazide functional group at the 5’ end of the stem-loop construct was coupled to the aldehyde functional group of the SWCNT-tethered FDBP *via* a hydrazide-aldehyde coupling reaction. To achieve this, the device was incubated in 100 µM of the stem-loop construct in Reaction Buffer (100 mM sodium acetate (NaOAc), pH=4.5) overnight (for ∼20 hours) followed by a 2-hour reduction in Reduction Buffer (10 mM NaHPO_4_/Na_2_H_2_PO_4_,, pH = 7.0 and 50 mM sodium cyanoborohydride (NaCNBH_3_)). The device was then rinsed twice with 500 µL of Recording Buffer at 50 °C to remove any non-specifically bound RNA. The smFET experimental data in this work derives from this single RNA molecule tethered to the smFET device.

The smFET experiments were performed in Recording Buffer. Current trajectories were recorded at varying temperatures (35, 40, 45, and 50 °C) for 5 minutes at a *V*_*lg*_ of 0.1 V and a *V*_*sd*_ of 25 mV. To achieve the desired temperature, the microfluidic chamber was incubated in the heating chamber of the smFET instrument for 15 min at the specified temperature before data collection was initiated. The smFET device was sampled at a time resolution of 40 µs using a 5.3 kHz anti-aliasing filter.

### Thermal melting experiments

To validate that our single SWCNT-tethered RNA molecule recapitulated the ensemble thermodynamic stability of the UUCG stem-loop, we performed thermal melting experiments of the h-SL-bio construct by monitoring the circular dichroism (CD) of the RNA at 280 nm as a function of temperature over a temperature window from 5 °C to 95 °C using a Chirascan V100 CD spectrophotometer. For this thermal melting experiment, a 10 µM sample of the h-SL-bio construct was prepared in Recording Buffer. Prior to the thermal melting experiment, the stem-loop construct was folded by pre-heating to 95 °C and subsequent cooling to 5 °C at a ramp rate of 2 °C min^−1^. CD measurements were recorded at every 0.2 °C increment with a ramp rate of 1 °C min^−1^ (Figure S3). Using a two-state model comprised of unimolecular transitions between single ***U*** and ***Z*** conformations [20], the equilibrium changes in enthalpy 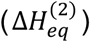 and entropy 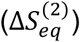 of folding for the stem-loop, where the superscript of (2) is used to denote the two-state model described above, was calculated using the following set of equations:

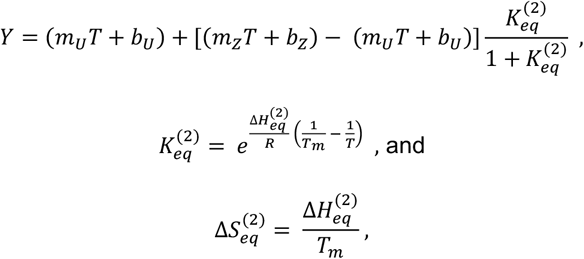

where *Y* is the observed CD signal, *m*_*U*_ and *m*_/_ are the baseline slopes for the ***U*** and ***Z*** states, *b*_*U*_ and *b*_/_ are the corresponding baseline intercepts, 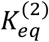 is the two-state equilibrium constant for the zipping reaction, *R* is the universal gas constant (1.987 cal K^−1^ mol^−1^), *T* is the absolute temperature of the measurement, and *T*_*m*_ is the absolute ‘melting’ temperature at which the corresponding free energy 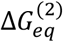 (given by 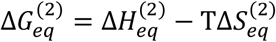) is zero. The above equations were fit to data collected for two independent replicates of the thermal melting experiment using a nonlinear curve fitting algorithm implemented with Matlab to yield the values of 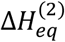 and 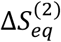 for the stem-loop (Table S2). These values were subsequently used to calculate the 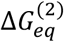 for the UUCG stem-loop at the temperatures for the smFET experiments (Figure S4).

An additional set of thermal melting experiments were performed for the h-SL construct in a manner identical to the h-SL-bio construct described above (Figure S3). The data from two replicates of this thermal melting experiment was similarly used to estimate the 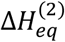 and 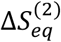 for the h-SL construct (Table S2).

### Identification of current states within current trajectories

The current trajectories for all of the experimental temperatures (35, 40, 45, and 50 °C; Figure S2) were first individually background corrected using an iterative detection algorithm (as previously described in [34, 36, 53]) to correct for the slowly drifting current baseline inherent to smFET devices [34, 36, 38]. The current trajectories at each temperature were subsequently divided into 200-ms time intervals, each containing 5,000 data points. Each interval was normalized such that the lowest current value for the interval corresponded to a normalized current signal of 0 a.u., and the highest current value corresponded to 1 a.u. This normalization further corrected for long-time-scale changes in the signal due to any remaining background current drift in the smFET device [36]. Intervals where excessive local background current drift and/or noise in the smFET device obscured transitions in the current trajectory resulting in an inability to clearly distinguish the two current states were discarded. In total, such intervals represented less than 3-10% of the total intervals collected at each temperature point.

At all temperatures other than 35 °C, the two current states present in the current trajectories could be identified and characterized using Gaussian mixture models (GMMs) inferred by employing a variational Bayesian expectation-maximization algorithm [54] implemented in the single-molecule data analysis platform, tMAVEN [55]. These GMMs yielded the means and standard deviations of the two observed states (Figure S2), which were subsequently used to generate idealized trajectories from the experimental data (based on the responsibilities [54] of the individual data points for each Gaussian distribution). The high-current state was centered around a normalized current value of ∼0.7 a.u. and the low-current state was centered around a normalized current value of ∼0.3 a.u. For the 35 °C dataset, the relative proportion of the low-current state was so small that the signal for this state could not be reliably described as a Gaussian distribution using a GMM. In this case, a threshold of 0.4 a.u. (which corresponded to a normalized current value three standard deviations below the mean of the high-current state) was utilized to differentiate between the two states. Based on a previous smFET study of RNA stem-loops, the high-current state was assigned to zipped (***Z***) conformations of the stem-loop, and the low-current state was assigned to unzipped (***U***) conformations [36].

### Inference of kinetic rates from current trajectories

Based on the idealized trajectories calculated as described above, dwell times in both ***U*** and ***Z*** conformations (defined as the contiguous time period the molecule spent in a ***U*** or ***Z*** conformation before transitioning to a ***Z*** or ***U*** conformation, respectively) were calculated. These dwell times were then used to generate the corresponding survival probability distributions, *F*(*t*), defined as the fraction of dwells that were greater than a specific *t*, for each conformation (Figures S5 and S6). Each survival probability distribution was fit to a single-exponential decay of the form

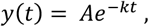

where *k* represents the rate constant and *A* is the pre-exponential term and a double-exponential decay of the form

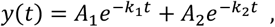

where *k*_1_ and *k*_2_ represent the rate constants for the two kinetic phases, and *A*_1_ and *A*_2_ are the respective pre-exponential terms for the two phases that also give their relative proportions. Subsequently, residuals of the fits,

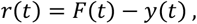

were also calculated. These residuals showed that the single-exponential fits missed the initial fast decay phase of the survival probability distribution for both conformations at all temperatures (Figures S5 and S6). Thus, the double-exponential decays better explained the survival distributions. The double-exponential decays each yielded a slow phase (*k*_*slow*_ and *A*_sl*o*w_) that arises from a long-lived RNA conformation (***U***^***L***^ or ***Z***^***L***^) and a fast phase (*k*_*fast*_ and *A*_*fast*_) that arises from a short-lived RNA conformation (***U***^***S***^ or ***Z***^***S***^). The kinetic parameter values we obtained from the double-exponential fits at all temperatures for the zipping and unzipping transitions are compiled in Tables S4 and S5, respectively.

### Calculation of activation enthalpies and entropies using transition state theory

Each *k* was converted to the corresponding activation free energy (Δ*G*^‡^) (Figure S7a-b) using the relation specified by transition state theory [48]

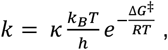

where *T* is the absolute temperature, *k*_*B*_ is the Boltzmann constant, *h* is the Planck constant, *R* is the universal gas constant, and *κ* is a correction factor that, in our calculations, is set to 1 [48]. For each kinetic phase in both the zipping and the unzipping transitions, the corresponding Δ*G*^‡^s over the experimental range of temperatures were used to estimate the Δ*H*^‡^s and Δ*S*^‡^s using a linear fit to

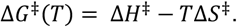

The Δ*H*^‡^ s and Δ*S*^‡^ s were assumed to be temperature-independent in the relatively small temperature range of the reported experiments, which agreed with the observed linear dependence of Δ*G*^‡^ on temperature (Figure S7a-b). The calculated Δ*G*^‡^s for both phases at all temperatures, and the fitted Δ*H*^‡^s and Δ*S*^‡^s, are compiled for the zipping and unzipping transitions in Table S6.

### Calculation of the equilibrium enthalpies and entropies between the kinetic phases

For both ***U*** and ***Z*** conformations, the relative proportions of the slow and fast kinetic phases were given by their respective pre-exponential terms (*A*_*slo*w_ and *A*_f*ast*_) in the double-exponential fits (Tables S4 and S5). Since these phases arose from the individual populations of the long-lived

(***U***^***L***^ or ***Z***^***L***^) and short-lived (***U***^***S***^ or ***Z***^***S***^) conformations, the ratios between these two terms 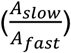 were interpreted as the ratios between the proportions of these long-lived and short-lived RNA conformations. This allowed us to calculate the equilibrium constants (*K*_*eq*_) between the two ***U*** conformations (and similarly, the two ***Z*** conformations) as

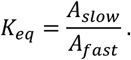

Subsequently, these *K*_*eq*_s, calculated over varying temperatures, were converted to the corresponding free energy differences, Δ*G*_*eq*_, at these temperatures as

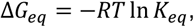

where *T* is the absolute temperature and *R* is the universal gas constant. Similar to the activation parameters, these Δ*G*_*eq*_s (Figure S7c-d) were fit to the linear equation

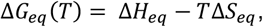

where Δ*H*_*eq*_ and Δ*S*_*eq*_ are the corresponding equilibrium enthalpic and entropic differences, that are also assumed to be temperature independent in the relatively small experimental range of temperatures studied here. The calculated Δ*G*_*eq*_s at all temperatures, and the fitted Δ*H*_*eq*_s and Δ*S*_*eq*_s, are compiled in Table S7.

## Acknowledgements

We thank Dr. Hashim al-Hashimi (Columbia University) and Dr. Jeffrey Kieft (New York Structural Biology Center) for valuable comments on the manuscript. We also thank Dr. Sarah Dubnik and other past and present members of the Gonzalez, Nuckolls, and Shephard labs for their assistance, comments, and advice. This work was carried out in part in the Clean Room, Electron Microscopy, and Shared Materials Characterization labs of Columbia Nano Initiative (CNI) Shared Lab Facilities and the Precision Biomolecular Characterization Facility (PBCF) at Columbia University.

## Authors Contributions

S.S.J., K.K.R., D.G.L., C.N., and R.L.G. designed the research; S.S.J. and D.G.L. fabricated the smFET devices; S.S.J. and D.G.L. performed the smFET experiments and collected the data; S.S.J. and K.K.R. analyzed the data; S.S.J., K.K.R., D.G.L., C.N., and R.L.G. wrote the manuscript; all authors approved the final version of the manuscript.

## Data and materials availability

The open-source Python code for tMAVEN, which was used for the single-molecule data analysis in this work, is freely available *via* a Git repository (https://github.com/GonzalezBiophysicsLab/tmaven). The current trajectories will be made freely available via Zenodo repository at the time of publication.

## SUPPORTING INFORMATION

### Conductance trajectories arise from the zipping dynamics of a single stem-loop

To validate that the observed conductance trajectories in our experiments arise from the zipping dynamics of a single, surface-tethered stem-loop RNA, we first determined the ensemble, two-state, differences in free energies, 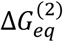, for zipping of the UUCG stem-loop, where the superscript denotes our assumption of the two-state model, using thermal melting experiments (Figure S3; see Materials and Methods). To determine the analogous 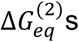 for zipping of the UUCG stem-loop using our smFET data, we applied a Gaussian mixture model (GMM)-based kinetic model [1] that identified two current states in our conductance trajectories. For the lowest temperature point of 35 °C, due to the very small population of the low-current state, a threshold was used, instead of a GMM, to distinguish between the two states (see Materials and Methods). Assuming that each state corresponded to a single RNA conformation, we were able to assign to the low- and high-current states to the unzipped ***U*** and the zipped ***Z*** conformations of the stem-loop (Figures 1 and S2). We note that the application of the GMM for the higher temperature points was made possible by an improved signal-to-noise ratio (SNR) of ∼4–5 in this work relative to the SNR of ∼3-4 that we have observed in our previous smFET studies of RNA stem-loops [2]. This improvement could be attributed to several changes in the experimental design of the present work *versus* our previous work, including the use of a lower ionic strength buffer, resulting in reduced screening of the electrostatic charges on the RNA [3, 4]; the use of an RNA stem-loop construct where the conformational changes induced by zipping and unzipping were more proximal to the surface of the single-walled carbon nanotube (SWCNT); and the direct, covalent functionalization of the RNA stem-loop construct to the surface of the SWCNT [5-7] (Figure S1). These improvements over our previous work increased the change in current through the SWCNT during the conformational rearrangements of the RNA molecule, resulting in the higher SNR observed in the present work.

Using our GMM- and threshold-based kinetic models, we calculated the fractional occupancies of the ***Z*** and ***U*** conformations (*f*_Z_ and *f*_*U*_, respectively) across the temperatures investigated here (Figure S2). In agreement with a previous study [2], *f*_*U*_ was observed to increase with increasing temperature, which is in line with the expectation that the unzipped ***U*** conformation should become more populated with increasing temperature (Table S3). These fractional occupancies were subsequently used to calculate the two-state equilibrium constants,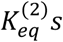, for the zipping reaction using

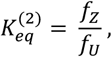

and the corresponding two-state equilibrium free energies of zipping using

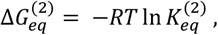

where *R* denotes the universal gas constant and *T* is the absolute temperature of the measurement. The 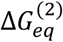 for the smFET experiments were in striking agreement with those from the ensemble thermal melting experiments (Figure S4a and Table S3), validating that the current trajectories obtained from our smFET experiments report on the same zipping reaction that the ensemble thermal melting experiments report on.

Next, using the 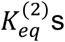 estimated from the thermal melting experiments, we calculated the theoretical fractional occupancies of the ***U*** conformation for a given number of surface-tethered stem-loops over the range of experimental temperatures investigated here (Figure S4b). In the case of a single RNA stem-loop tethered to a single smFET device, *f*_*U*_ was directly calculated from 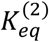 as

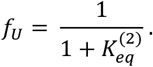

For the case of multiple RNA stem-loops tethered to a single smFET device, we assumed that the ***U*** conformation corresponded to the scenario where at least one stem-loop was unzipped and the ***Z*** conformation corresponded to the case where all stem-loops were zipped. Given these assumptions, the expected fractional occupancy of the ***U*** conformation for this case was given by

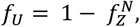

where *N* denotes the number of tethered stem-loops and *f*_&_ is the fractional occupancy of a single RNA stem-loop at a given temperature, which can be calculated from the ensemble 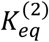 using

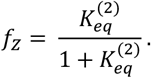

Comparing the *f*_*U*_ values obtained from the GMM-based kinetic modeling employed to analyze for our experimentally obtained current trajectories with the *f*_*U*_ values that would be expected for various different numbers of surface-tethered RNA stem-loops allowed direct determination of the number of RNA stem-loops that generated a particular, experimentally obtained current trajectory. The results of this comparison showed that the *f*_*U*_ values we experimentally obtained overwhelmingly agree with the *f*_*U*_ values expected for the case in which a single RNA stem-loop is tethered to the surface of the SWCNT, confirming that our experimental conductance trajectories arose from a single, surface-tethered RNA molecule (Figure S4b). We note that the analysis described here may be used in the future as a general framework for directly determining the number of surface-tethered biomolecules on any smFET device.

### Identification of the kinetic scheme between observed RNA conformations

To determine the kinetic scheme governing the zipping dynamics of the UUCG stem-loop, we made use of the fact that equilibrium changes in enthalpy and entropy (Δ*H*_*eq*_ and Δ*S*_*eq*_, respectively) are thermodynamic state variables. Thus, for any two given states, these quantities should depend only on the identities of the states and not on the paths used to calculate them. We considered the two ***U*** conformations (***U***^***L***^ and ***U***^***S***^) and two different paths connecting them, the ‘direct’ path A and the ‘indirect’ path B (Figures 2 and S7). For path A, Δ*H*_*eq*_(***U***^***L***^, ***U***^***S***^) and Δ*S*_*eq*_(***U***^***L***^, ***U***^***S***^) between these conformations, where ***U***^***S***^ represents the initial conformation and ***U***^***L***^ the final conformation, were calculated directly from the proportion of the slow and fast kinetic phases that corresponded to these ***U*** conformations (Figure S8c-d and Table S7; see Materials and Methods).

The indirect path B involves the two ***Z*** conformations (***Z***^***L***^ and ***Z***^***S***^) (Figures 2 and S7). In this path, Δ*H*_*eq*_(***U***^***L***^, ***U***^***S***^) and Δ*S*_*eq*_(***U***^***L***^, ***U***^***S***^) can be expressed as differences between the ***U*** conformations and the ***Z*** conformations, and between the ***Z*** conformations themselves [8]. As above, Δ*H*_*eq*_(***Z***^***L***^, ***Z***^***S***^) and Δ*S*_*eq*_(***Z***^***L***^, ***Z***^***S***^) can be directly calculated based on the proportion of the slow and fast kinetic phases that corresponded to these ***Z*** conformations (Figure S8c-d and Table S7; see Materials and Methods). However, the Δ*H*_*eq*_ s (and Δ*S*_*eq*_ s) between the ***U*** and ***Z*** conformations cannot be calculated in a similar manner, because our dwell-time analysis (Figures S5 and S6) only yielded the rates of transitions out of a particular conformation and not the rates between two conformations [9]. Instead, these rates yielded the activation parameters (Δ*H*^‡^s and Δ*S*^‡^) for transitions out of these conformations (Figure S8a-b and Table S6; see Materials and Methods). These Δ*H*^‡^ s and Δ*S*^‡^ represented the corresponding changes between the ***U***, and between the ***Z***, conformations and the transition states (***TS***^‡^s) for the transitions exiting these conformations [10]. To calculate the corresponding Δ*H*_*eq*_s and Δ*S*_*eq*_s, we need to first specify how these ***TS***^‡^s fit into the kinetic scheme.

For kinetic scheme 1 (Figures 2a and S7c), we assume that the ***TS***^‡^ exiting out of ***U***^***S***^ 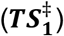 corresponds to that out of ***Z***^***S***^. We can then estimate the Δ*H*_*eq*_ and Δ*S*_*eq*_ between them as

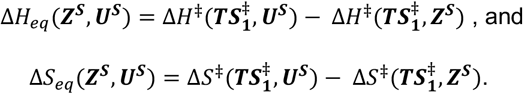

In this case, the other 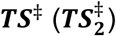 out of ***U***^***L***^ would correspond to that out of ***Z***^***L***^, resulting in

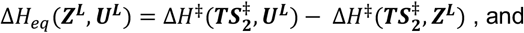

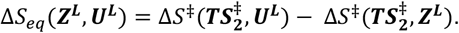

Thus, the ‘indirect’ Δ*H*_*eq*_(***U***^***L***^, ***U***^***S***^) and Δ*S*_*eq*_(***U***^***L***^, ***U***^***S***^) can be written as

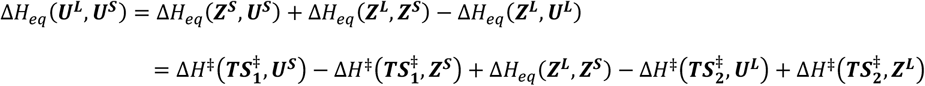

and

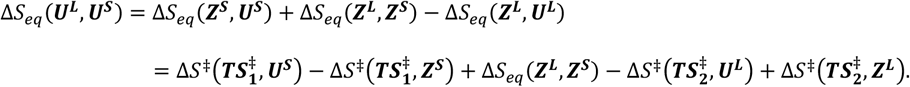

In this scheme, by placing the ***TS***^‡^s in the above manner, we have assumed that transitions only occur between ***U***^***S***^ and ***Z***^***S***^, and between ***U***^***L***^ and ***Z***^***L***^ (Figures 2a and S7c).

Alternatively, in kinetic scheme 2 (Figures 2b and S7d), we assume that the ***TS***^‡^ exiting out of 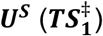 corresponds to that out of ***Z***^***L***^, resulting in

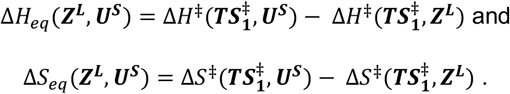

In this case, the other ***TS***^‡^ out of 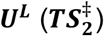 would correspond to that out of ***Z***^***S***^, resulting in

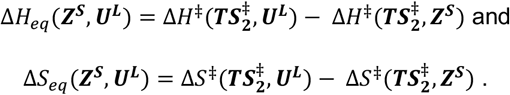

Thus, in this case, the ‘indirect’ Δ*H*_*eq*_ and Δ*S*_*eq*_ between ***U***^***L***^ and ***U***^***S***^ can be written as

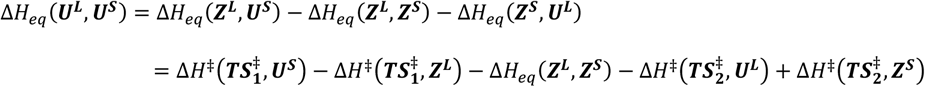

and

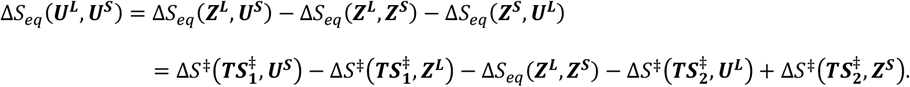

In this scheme, with the above placement of the ***TS***^‡^s, we have assumed that transitions only occur between ***U***^***S***^ and ***Z***^***L***^, and between ***U***^***L***^ and ***Z***^***S***^ (Figures 2b and S7d).

While the two schemes differ only in how the ***TS***^‡^s, and thus the corresponding activation parameters, are arranged, we see that they lead to drastically different expressions for the indirect Δ*H*_*eq*_ and Δ*S*_*eq*_ between ***U***^***S***^ and ***U***^***L***^. By analyzing our experimentally obtained data in this way, we show that the zipping dynamics of the UUCG stem-loop are predominantly governed by kinetic scheme 1 (Figures 2 and S7c-d, right-most panels).

---

**Figure S1.**
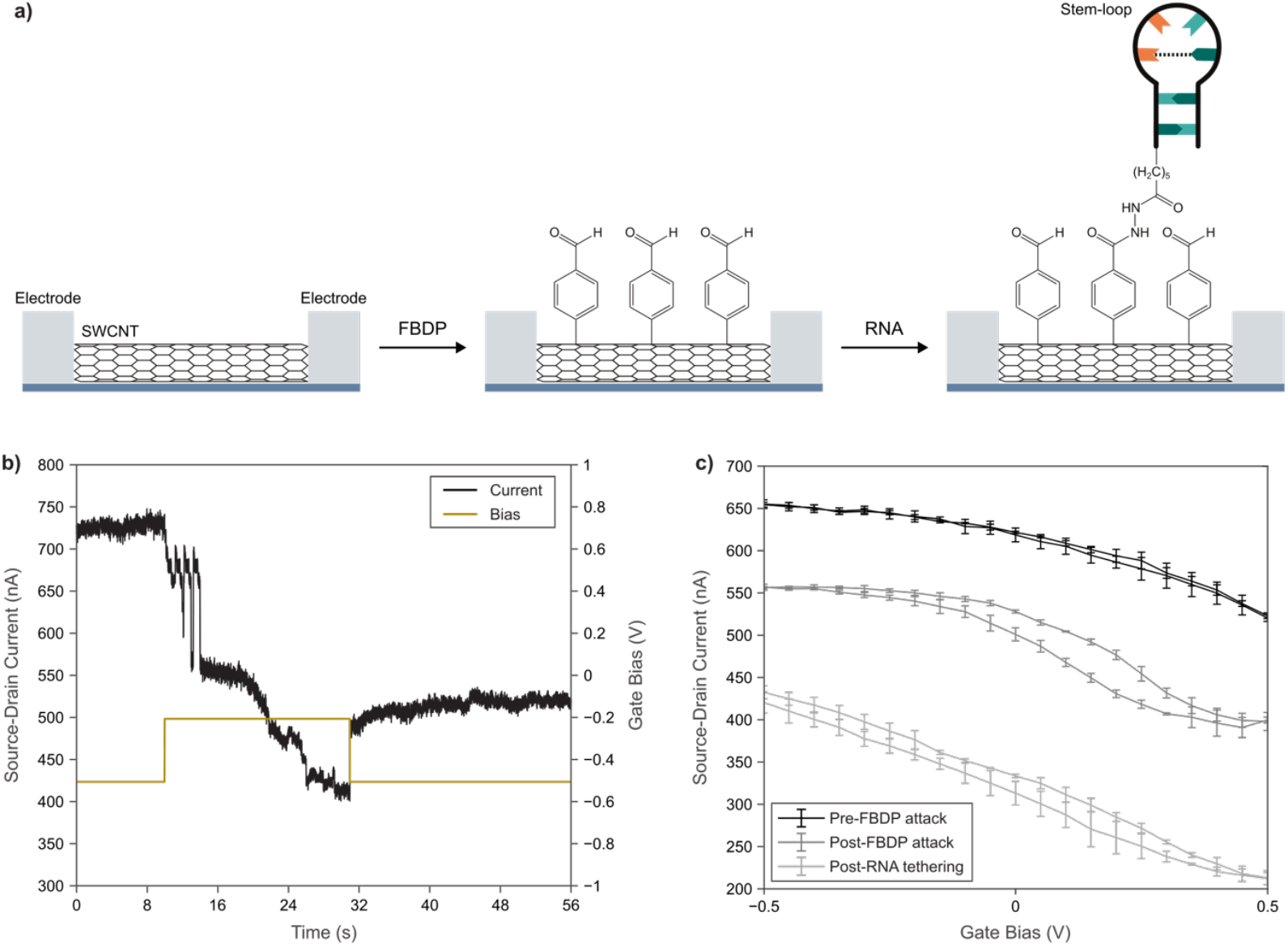
Covalent tethering of an RNA molecule to the surface of a SWCNT using an electrically controlled diazonium coupling reaction. **(a)** Schematic of the covalent tethering approach, showing first the tethering of FBDP onto the SWCNT surface through an electrically controlled diazonium reaction, followed by an RNA tethering step through a aldehyde-hydrazide coupling reaction. **(b)** *I*_*sd*_-*t* (black) and *V*_*lg*_-*t* (brown) measurements of an smFET device during the FBDP attack, showing multiple, clearly distinguished drops in current indicative of FBDP tethering when *V*_*lg*_ is ramped to −0.2 V from −0.5 V. **(c)** Average *I*_*sd*_-*V*_*lg*_ curves for the same smFET device before FBDP attack, after FBDP attack, and after RNA tethering, measured in Recording Buffer. Error bars denote the standard deviation of three measurements.

**Figure S2.**
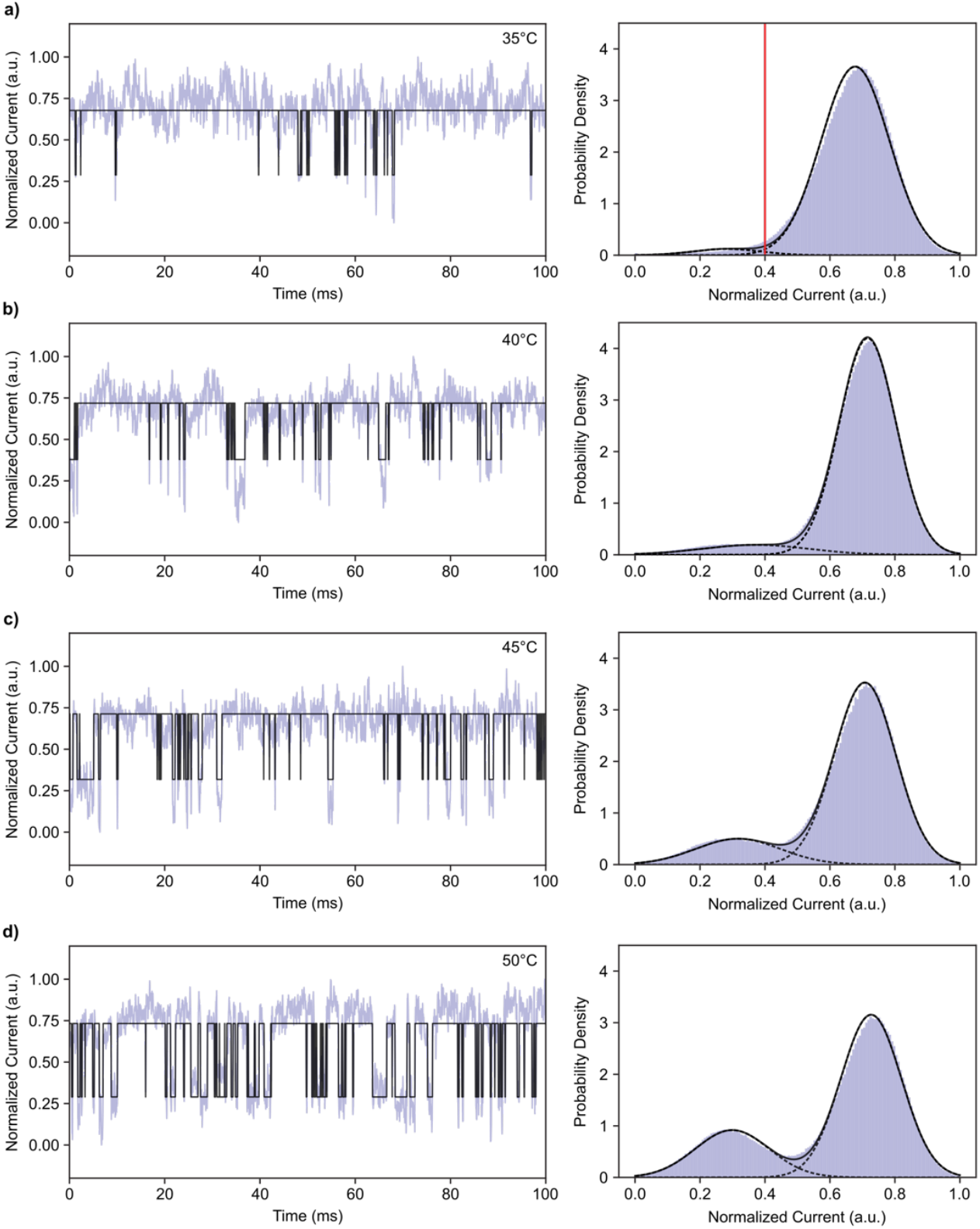
Temperature dependence of current trajectories recorded using the same, single UUCG stem-loop tethered onto the SWCNT surface of the same, single smFET device. smFET datasets obtained at **(a)** 35 °C, **(b)** 40 °C, **(c)** 45 °C, and **(d)** 50 °C showing (*left*) a representative 100-ms interval of the current trajectory (violet) overlaid with the idealized trajectory (black) and (*right*) a histogram of the current values across an entire 60 s trajectory (violet) overlaid with the inferred Gaussian density for each current state (dashed black curves) and the sum of all states (solid black curve) for the dataset at each temperature. The kinetic model that yielded these Gaussian distributions at 40 °C, 45 °C, and 50 °C were GMMs that were inferred from the data. For the 35 °C dataset, current values were first separated by a threshold value at 0.4 a.u. (red) and the means and standard deviations of the resulting populations yielded the individual Gaussian distributions for each state.

**Figure S3.**
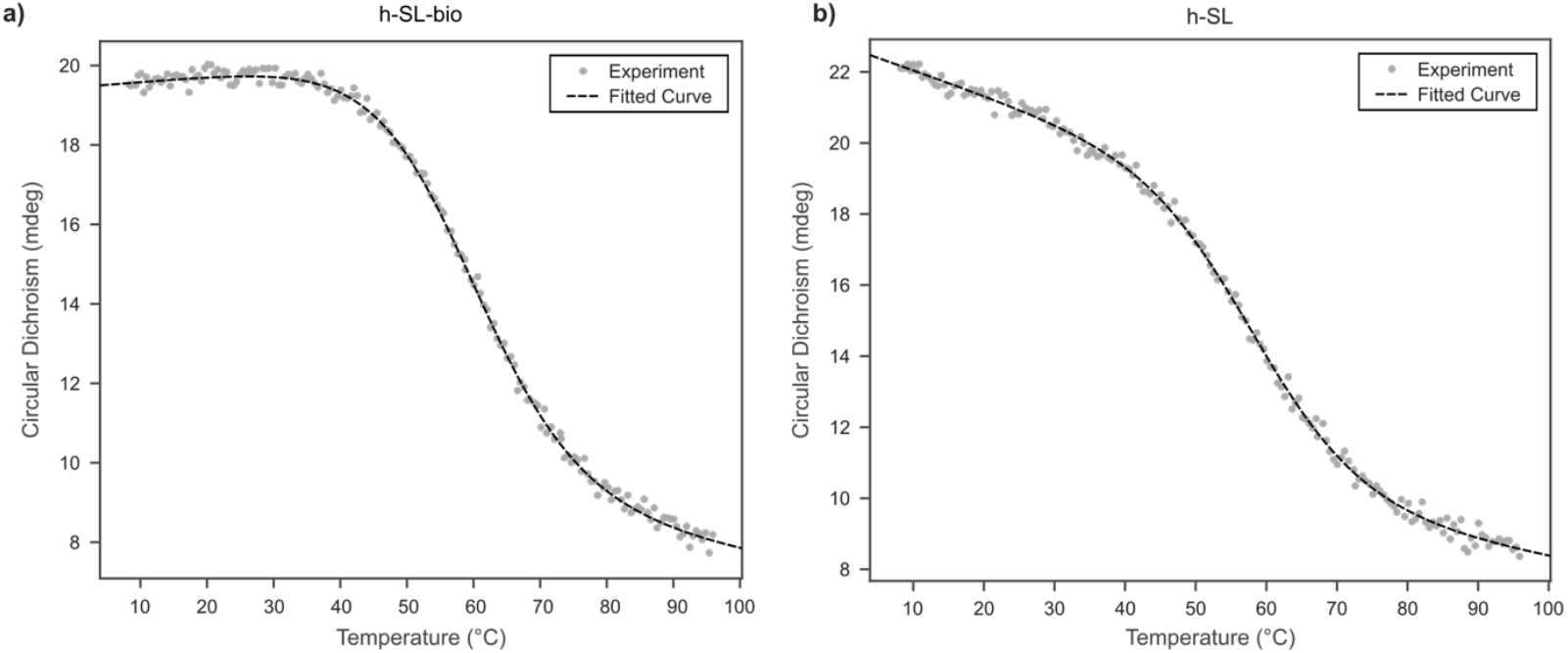
Thermal melting of the h-SL-bio and h-SL RNA stem-loop constructs. Thermal melting curves showing the CD measured at 280 nm (gray circles) for the **(a)** h-SL-bio and **(b)** h-SL constructs as a function of temperature, along with the fit (black curve) to a two-state model of unimolecular transitions between the ***U*** and ***Z*** conformations.

**Figure S4.**
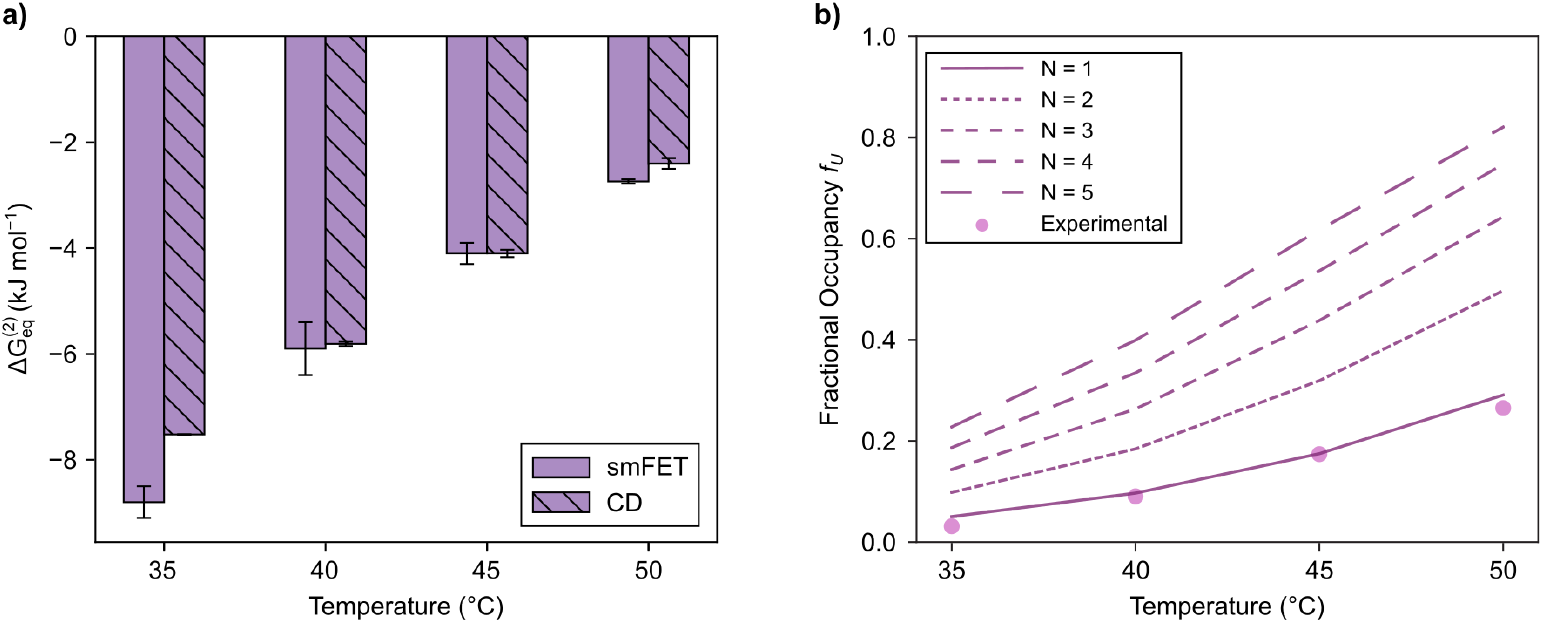
Confirmation that conductance trajectories report on the zipping dynamics of single, surface-tethered RNA stem-loops. **(a)** Comparisons of the 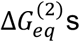 of zipping for the smFET (solid) and CD thermal melting (hatched) experiments. Error bars for the smFET experiments represent the standard deviations of the 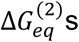 calculated from three separate 20-s windows for the current trajectories at each temperature. Error bars for the CD thermal melting experiments represent the standard deviations of the 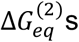 calculated from two independent experiments. **(b)** The expected fractional occupancies of the ***U*** conformation (*f*_*U*_) at each temperature for a single tethered RNA stem-loop and various multiple tethered RNA stem-loop (lines) along with the *f*_*U*_ calculated from experimental smFET data (circles).

**Figure S5.**
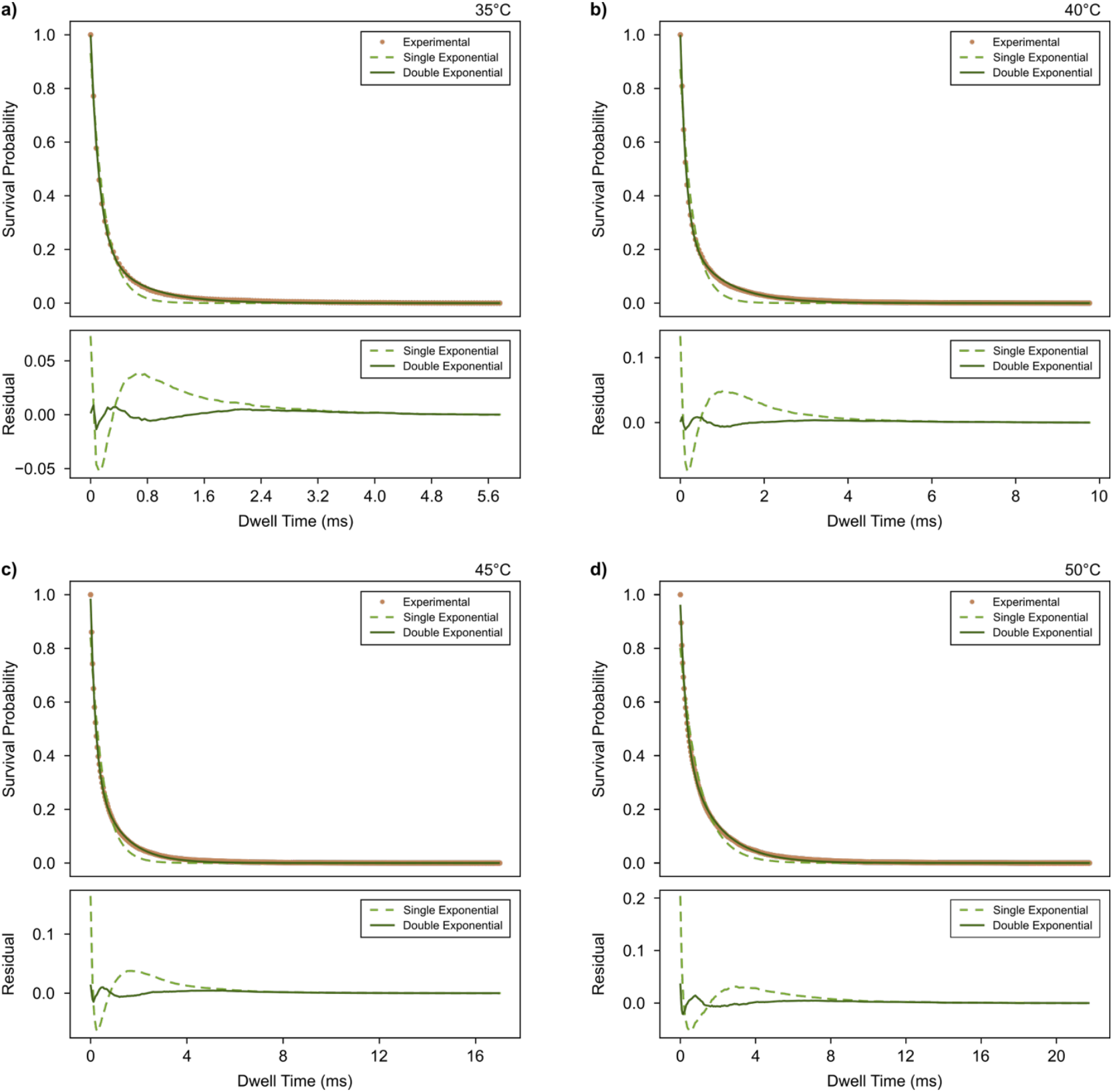
Kinetics of zipping. (*top*) Survival probabilities for the dwell-time distributions of the ***U*** conformation (circle) along with the single-exponential (dashed) and double-exponential (solid) fits and (*bottom*) the residuals of the single-exponential (dashed) and double-exponential (solid) fits for smFET datasets collected at **(a)** 35 °C, **(b)** 40 °C, **(c)** 45 °C, and **(d)** 50 °C.

**Figure S6.**
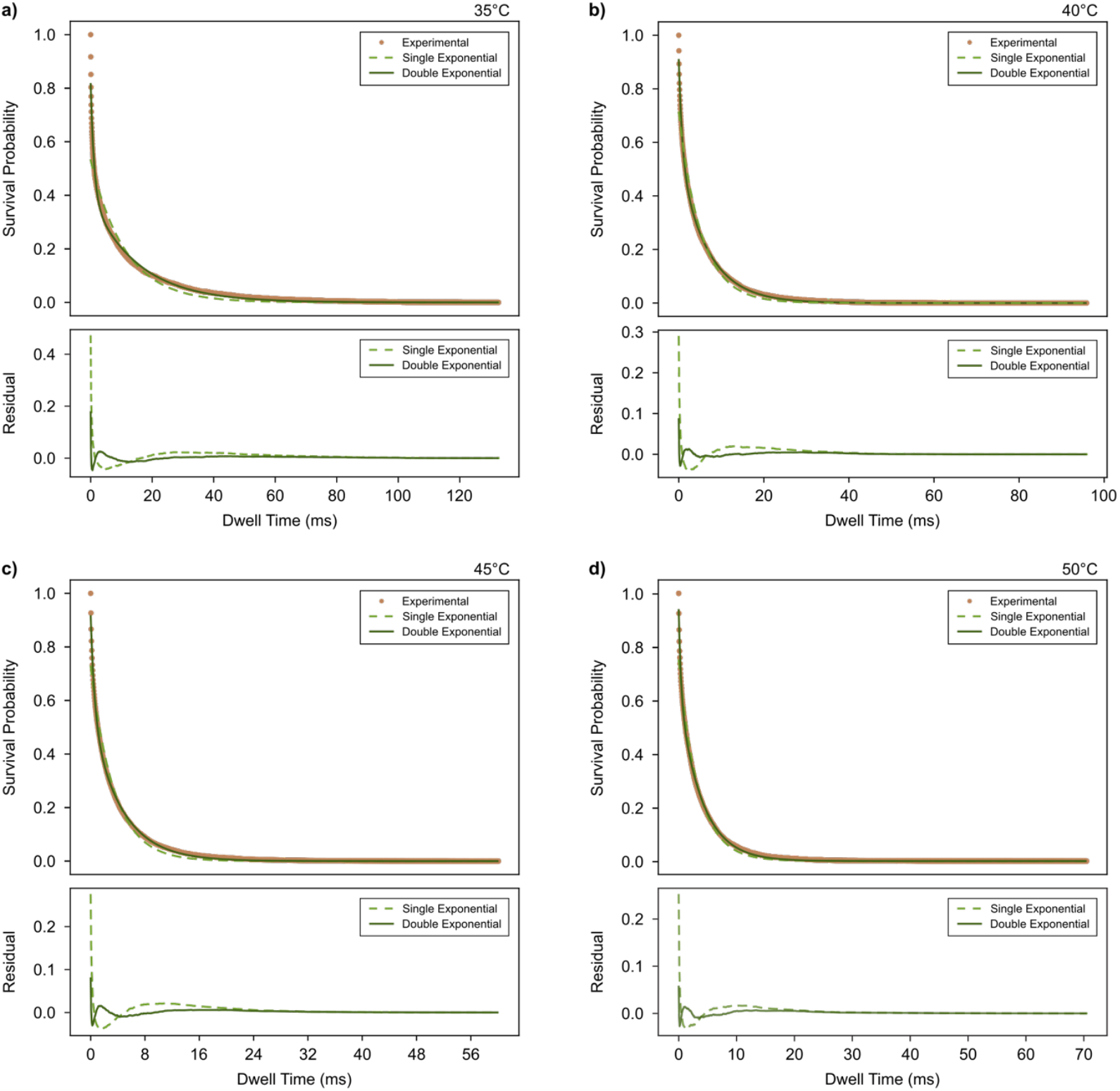
Kinetics of unzipping. (*top*) Survival probabilities for the dwell-time distributions of the ***Z*** conformation (circle) along with the single-exponential (dashed) and double-exponential (solid) fits and (*bottom*) the residuals of the single-exponential (dashed) and double-exponential (solid) fits for smFET datasets collected at **(a)** 35 °C, **(b)** 40 °C, **(c)** 45 °C, and **(d)** 50 °C.

**Figure S7.**
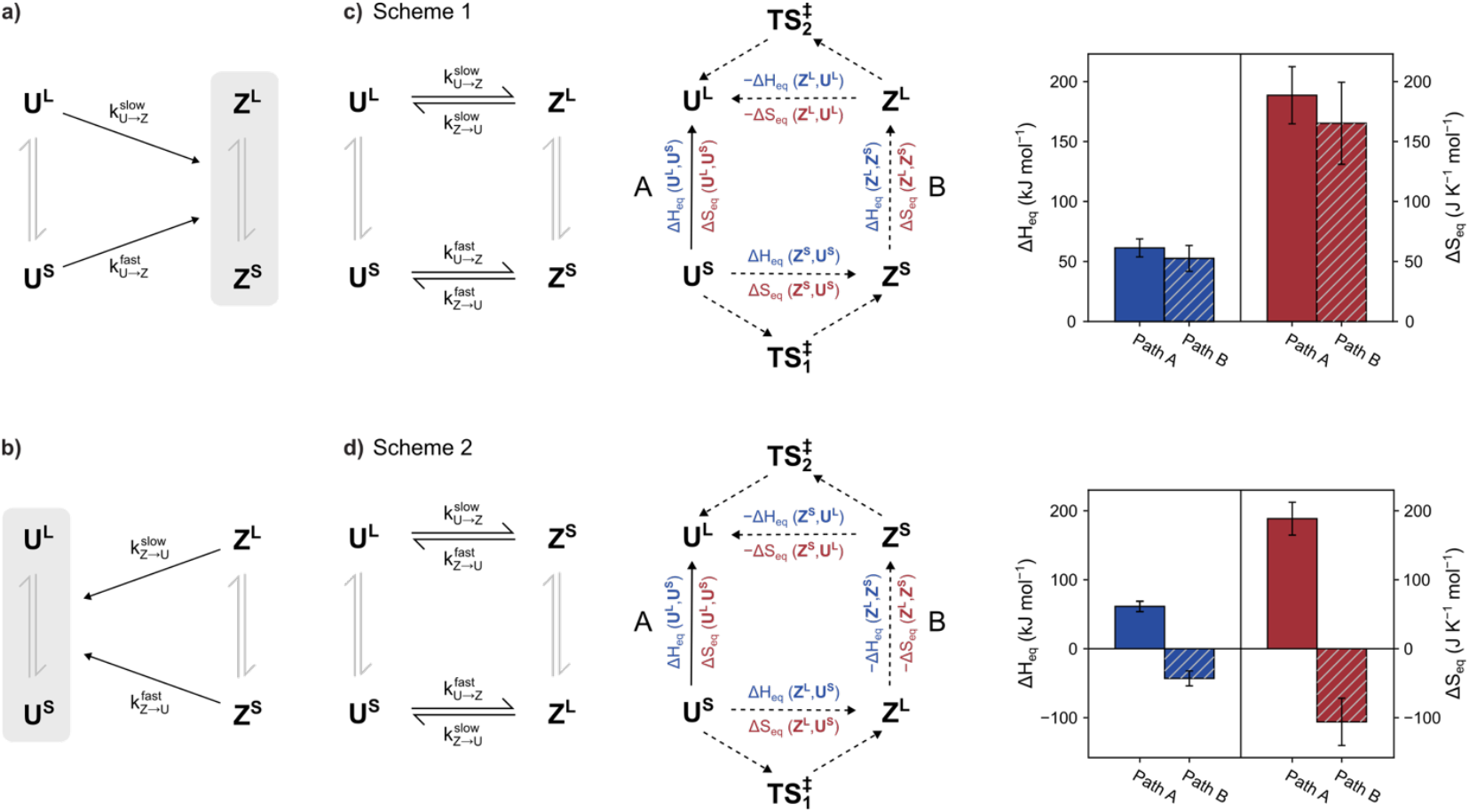
Potential kinetic schemes governing the zipping dynamics of the UUCG RNA stem-loop. The observed slow and fast kinetic phases, and the corresponding conformations that they transition from, for **(a)** zipping and **(b)** unzipping. **(c)** Potential kinetic scheme 1, showing *(left)* the predominant transitions between ***U***^***S***^ and ***Z***^***S***^ and between ***U***^***L***^ *and* ***Z***^***L***^, *(center)* the corresponding thermodynamic cycle depicting the direct path A (solid arrow) and the indirect path B (dashed arrows) between ***U***^***S***^ and ***U***^***L***^, and *(right)* the calculated Δ*H*_*eq*_*s* (blue) and Δ*S*_*eq*_*s* (red) for these paths. The unobserved transitions between the ***U*** and ***Z*** conformations are shown as grey equilibrium arrows. **(d)** Potential kinetic scheme 2, showing *(left*) the predominant transitions between ***U***^***S***^ *and* ***Z***^***L***^ and between ***U***^***L***^ and ***Z***^***S***^, *(center)* the corresponding thermodynamic cycle depicting the direct path A (solid arrow) and the indirect path B (dashed arrows) between ***U***^***S***^ and ***U***^***L***^, and *(right)* the calculated Δ*H*_*eq*_ *s* (blue) and Δ*S*_*eq*_ *s* (red) for these paths. The unobserved transitions between the ***U*** and ***Z*** conformations are shown as grey equilibrium arrows.

**Figure S8.**
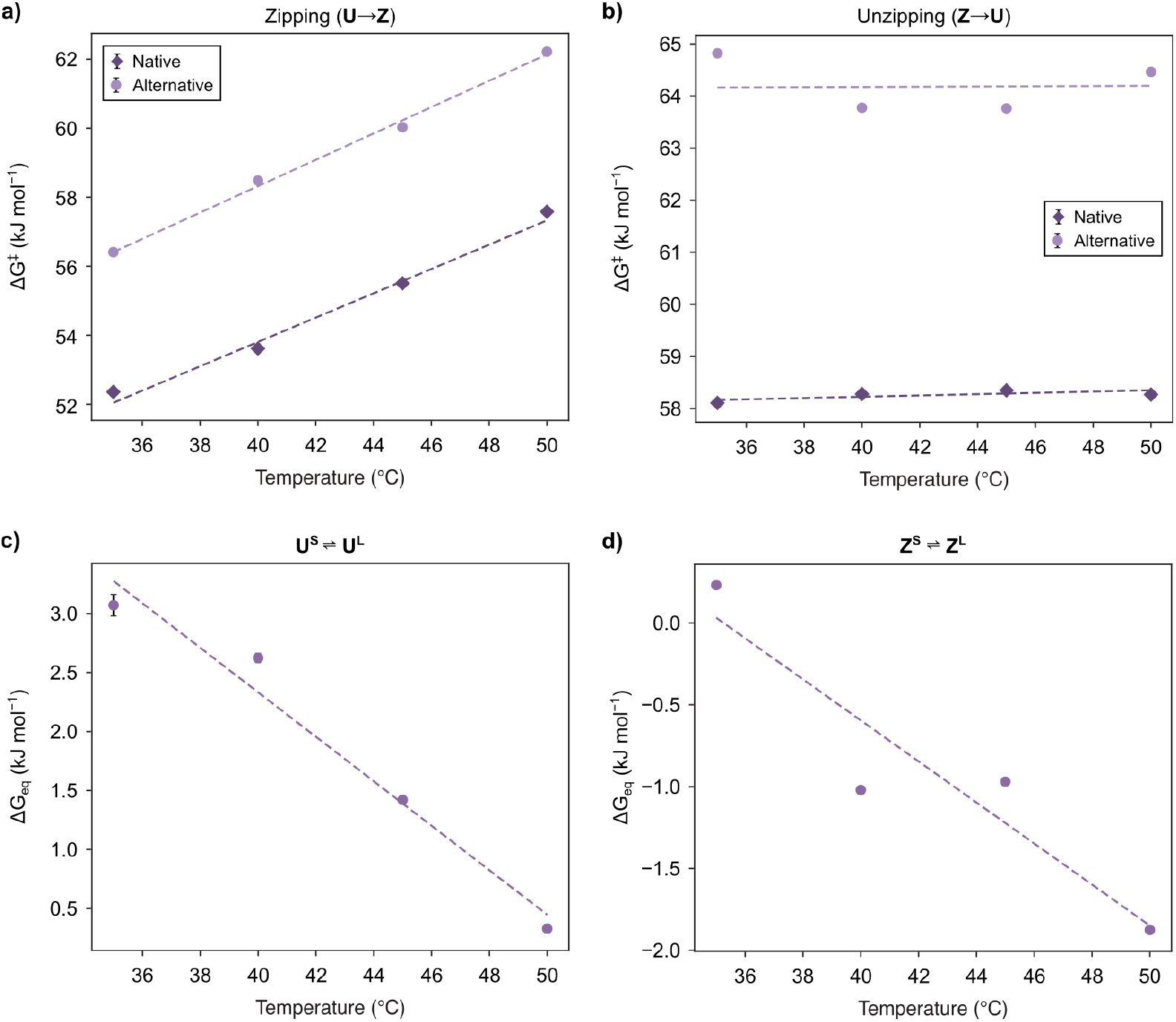
The activation and equilibrium free energies for the *U* and *Z*conformations. **(a)** Activation free energy of zipping (Δ*G*^‡^) as a function of temperature for the native (dark purple diamonds) and alternative (light purple circles) pathways along with linear fits to the data (dashed dark purple and dashed light purple lines, respectively). **(b)** Activation free energy of unzipping (Δ*G*^‡^) as a function of temperature for the native (dark purple diamonds) and alternative (light purple circles) pathways along with linear fits to the data (dashed dark purple and dashed light purple lines, respectively). **(c)** Equilibrium free energy (Δ*G*_*eq*_) for the ***U***^***S***^ ⇌ ***U***^***L***^ equilibrium as a function of temperature (circles) along with a linear fit to the data (dashed line). **(d)** Equilibrium free energy (Δ*G*_*eq*_) for the ***Z***^***S***^ ⇌ ***Z***^***L***^ equilibrium as a function of temperature (circles) along with a linear fit to the data (dashed line). Error bars in all plots represent the propagated error of the fit of the double-exponential decay function to the survival probability of the dwell-time distributions.

**Table S1.** Sequences of the RNA stem-loop constructs.

| **Stem-loop Construct** | **Sequence** |
| --- | --- |
| h-SL-bio | 5'-hydrazide-GC<u>UUC</u>GGC-biotin-3' |
| h-SL | 5'-hydrazide-GC<u>UUC</u>GGC-3' |

**Table S2.** Thermodynamic parameters for the two-state model of RNA stem-loop zipping based on CD thermal melting experiments. Parameters from the fit of the set of parametric equations for two-state, unimolecular transitions between the ***U*** and ***Z*** conformations to the data from the CD thermal melting experiments. The parameters are the average of fitting results for two independently prepared samples and the errors are the standard deviations of the fitting results.

| | $\Delta H_{eq}^{(2)}$<br>(kJ mol<sup>-1</sup>) | $\Delta S_{eq}^{(2)}$<br>(J mol<sup>-1</sup>K<sup>-1</sup>) | $\Delta G_{eq}^{(2)}(37)$<br>(kJ mol<sup>-1</sup>) | $T_m$<br>(°C) |
| --- | --- | --- | --- | --- |
| **h-SL-bio** | -113(2) | -342(8) | -6.84(0.01) | 57.0(0.5) |
| **h-SL** | -115.7(0.2) | -351(1) | -7.0(0.2) | 56.9(0.6) |

**Table S3.** Comparison of thermodynamic parameters determined from smFET and CD thermal melting experiments. Errors (in parentheses) for the parameters obtained from the analysis of the smFET experiments represent the standard deviations of the parameters calculated from three separate 20-s windows for the current trajectories at each temperature. Error (in parentheses) for the parameters obtained from the analysis of the CD thermal melting experiments represent the standard deviations of the parameters calculated from two independent experiments.

| | $\Delta G_{eq}^{(2)} \text{ (kJ mol}^{-1}\text{)}$ | | $f_u$ | |
| --- | --- | --- | --- | --- |
|  | smFET | CD | smFET | CD |
| **35°C** | -8.8(0.3) | -7.520(0.003) | 0.031(0.004) | 0.0504(0.0007) |
| **40°C** | -5.9(0.5) | -5.81(0.04) | 0.09(0.02) | 0.097(0.001) |
| **45°C** | -4.1(0.2) | -4.10(0.07) | 0.17(0.01) | 0.175(0.004) |
| **50°C** | -2.74(0.04) | -2.4(0.1) | 0.265(0.003) | 0.291(0.009) |

**Table S4.** The two kinetic phases describing the zipping of the UUCG stem-loop. Errors (in parentheses) represent the error of the fit for the double-exponential decay function to the survival probability of the dwell-time distributions of the ***U*** conformations at each temperature.

|  | Slow phase |  | Fast phase |  |
| --- | --- | --- | --- | --- |
| | $k_{U \rightarrow Z}^{slow} \text{ (s}^{-1}\text{)}$ | $A_{U \rightarrow Z}^{slow}$ | $k_{U \rightarrow Z}^{fast} \text{ (s}^{-1}\text{)}$ | $A_{U \rightarrow Z}^{fast}$ |
| **35°C** | 1740(40) | 0.231(0.008) | 8500(100) | 0.767(0.008) |
| **40°C** | 1130(10) | 0.267(0.004) | 7360(70) | 0.732(0.004) |
| **45°C** | 914(8) | 0.363(0.004) | 5060(60) | 0.622(0.004) |
| **50°C** | 584(4) | 0.452(0.005) | 3280(50) | 0.511(0.004) |

**Table S5.** The two kinetic phases describing the unzipping of the UUCG stem-loop. Errors (in parentheses) represent the error of the fit of the double-exponential decay function to the survival probability of the dwell-time distributions of the ***Z*** conformations at each temperature.

|  | Slow phase |  | Fast phase |  |
| --- | --- | --- | --- | --- |
| | $k_{Z \rightarrow U}^{slow} \text{ (s}^{-1}\text{)}$ | $A_{Z \rightarrow U}^{slow}$ | $k_{Z \rightarrow U}^{fast} \text{ (s}^{-1}\text{)}$ | $A_{Z \rightarrow U}^{fast}$ |
| **35°C** | 65.2(0.3) | 0.391(0.001) | 900(10) | 0.428(0.003) |
| **40°C** | 148.2(0.4) | 0.544(0.002) | 1220(10) | 0.367(0.002) |
| **45°C** | 222.5(0.9) | 0.542(0.003) | 1720(30) | 0.376(0.003) |
| **50°C** | 252.9(0.7) | 0.629(0.002) | 2540(40) | 0.313(0.003) |

**Table S6.**
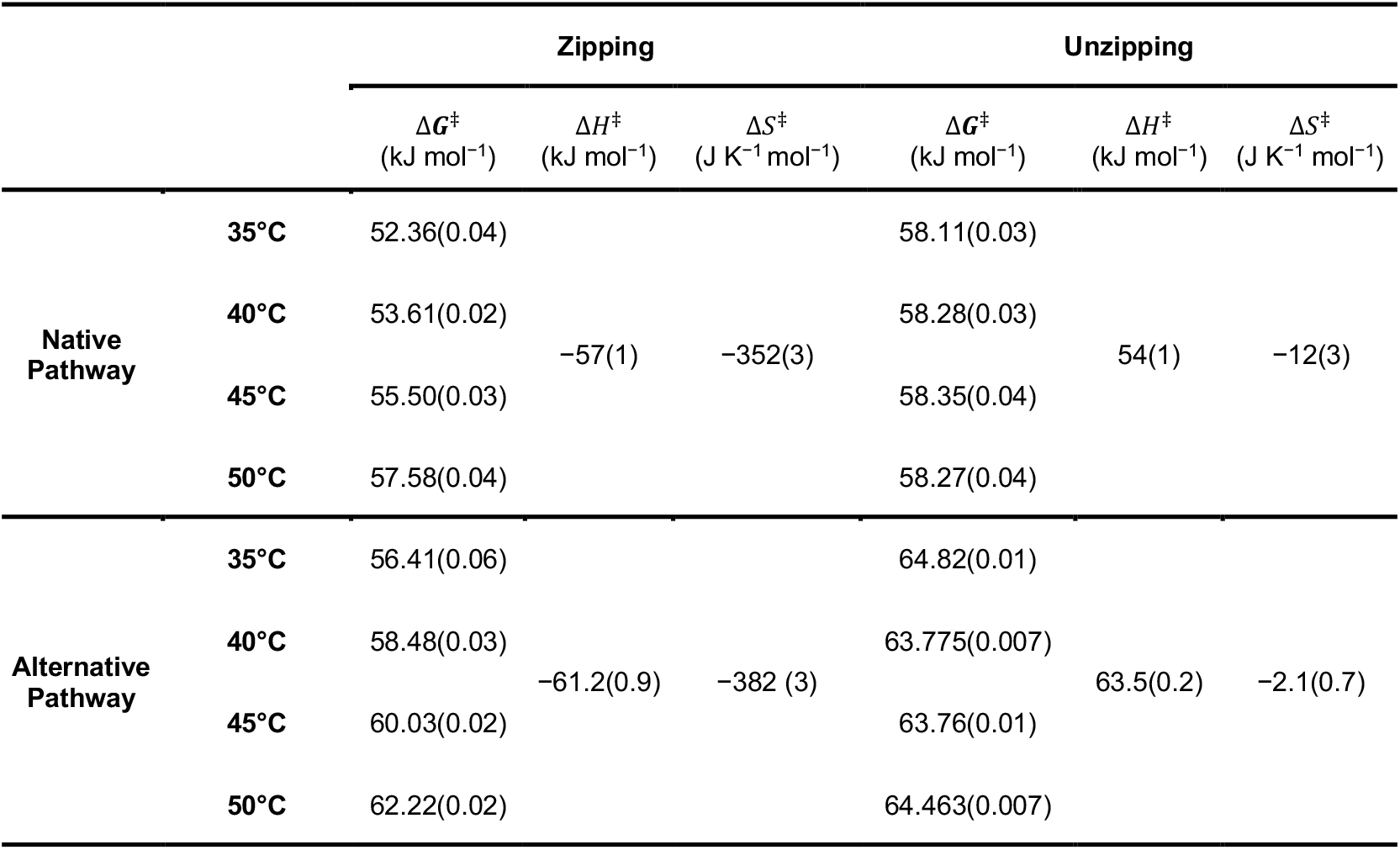
Activation free energies, enthalpies, and entropies of the zipping and unzipping of the UUCG stem-loop along the native and alternative pathways. Errors (in parentheses) of the Δ*G*^‡^s are propagated from the corresponding transition rate constants. Errors for Δ*H*^‡^s and Δ*S*^‡^s represent the error of the fit of the linear function to the Δ*G*^‡^s as a function of absolute temperature.

**Table S7.** Equilibrium free energies, enthalpies, and entropies for the two kinetic phases of zipping and unzipping of the UUCG stem-loop. Errors (in parentheses) of the Δ*G*_*eq*_s are propagated from the corresponding ratio of pre-factors for the double-exponential decay. Errors for Δ*H*_*eq*_s and Δ*S*_*eq*_s represent the error of the fit of the linear function to the Δ*G*_*eq*_s as a function of absolute temperature.

| | | $\Delta G_{eq}$<br>(kJ mol<sup>-1</sup>) | $\Delta H_{eq}$<br>(kJ mol<sup>-1</sup>) | $\Delta S_{eq}$<br>(J K<sup>-1</sup> mol<sup>-1</sup>) |
| --- | --- | --- | --- | --- |
| $U^S \rightleftharpoons U^L$ | 35°C | 3.07(0.09) | 61(8) | 190(20) |
|  | 40°C | 2.62(0.04) |  |  |
|  | 45°C | 1.42(0.04) |  |  |
|  | 50°C | 0.33(0.04) |  |  |
| $Z^S \rightleftharpoons Z^L$ | 35°C | 0.23(0.02) | 40(10) | 130(30) |
|  | 40°C | -1.02(0.02) |  |  |
|  | 45°C | -0.97(0.02) |  |  |
|  | 50°C | -1.88(0.02) |  |  |

